# A selective Cullin 3 RING E3 ligase inhibitor attenuates hyperglycemia via dual insulin sensitizing and insulinotropic action

**DOI:** 10.64898/2026.01.28.702366

**Authors:** Lijie Gu, Lei Xiong, Mohammad Nazmul Hasan, Yanhong Du, Timothy Wu, Tiangang Li

## Abstract

Hyperglycemia is a hallmark of type-2 diabetes and a key pathogenic driver of diabetic complications. Cullin RING E3 ligases (CRLs) are multi-subunit E3 ubiquitin ligases that mediate cellular protein turnover. The activity of CRLs requires cullin neddylation, a post-translational modification that can be pharmacologically targeted with therapeutic potentials. By using hyperinsulinemic-euglycemic clamp analysis, we discover that pan neddylation inhibitor exerts both insulin sensitization effect in liver and muscle and insulinotropic effect in pancreatic β cells. This dual action is mediated by Cullin 3 (Cul3), a member of the 7 canonical cullin family proteins. DI-1859, a selective Cul3 neddylation inhibitor, effectively protects against hyperglycemia in obese mice. DI-1859 enhances insulin signaling by preventing Cul3-mediated insulin receptor substrate degradation in liver and muscle cells. DI-1859 increases insulin secretion in a glucagon-like peptide-1-independent manner in mice and directly potentiates glucose-stimulated insulin secretion in INS-1 832/13 β cells and human islets. Mechanistic studies reveal that DI-1859 does not promote glycolytic flux or bioenergetics function but potentiates glucose-stimulated insulin secretion via mechanisms involving RhoA activation and cytoskeleton remodeling in β cells. This study shows that a single agent targeting Cul3 neddylation simultaneously promotes insulin sensitization and insulin secretion to attenuate hyperglycemia in mice.

**Article Highlights:**

a. Pan cullin neddylation inhibitors exhibit potent hypoglycemic effect.
b. The target organs and mechanisms underlying the hypoglycemia effect of cullin pan neddylation inhibitors are incompletely understood.
c. We found that inhibition of Cul3 leads to a dual insulin sensitization and insulinotropic effect.
d. Selective inhibition of Cul3 neddylation is a feasible approach to lower hyperglycemia.

## Introduction

Cullin RING E3 ligases (CRLs) are a sub-class of ubiquitin E3 ligases (1). A functional CRL is a multi-protein complex consisting of a cullin scaffold protein that interacts with a substrate receptor that recruits target proteins and a RING E3 ligase that ubiquitinates the target proteins for subsequent proteasomal degradation (2). Mammalian cells express 7 canonical cullin proteins, each of which can interact with a unique set of substrate receptors in a tissue-specific manner. It is estimated that CRLs may be responsible for ∼20% of the total cellular protein turnover, through which they regulate diverse cellular pathways and function (1, 3). An important regulatory feature is that CRLs are only functionally active upon cullin neddylation, an enzymatic reaction that covalently conjugates a ubiquitin-like protein called Nedd8 to a conserved lysine on cullin protein (2). Neddylation is mediated by a set of Nedd8 E1, E2 and E3 enzymes that are highly selective toward cullin proteins (4). The Nedd8-activating E1 enzyme (NAE1) is the only known mammalian Nedd8 E1 enzyme and its inhibition by MLN4924 and TAS4464 decreases the activity of all CRLs (2). In recent years, CRLs have been recognized as promising therapeutic targets for cancer therapy due to their roles in regulating cell proliferation and survival (5).

Hyperglycemia is a key cause of macrovascular and microvascular complications of type-2 diabetes (6). Improving hepatic, muscle and adipose insulin sensitivity and promoting β cell insulin secretion are the major therapeutic strategies for glycemic control in type-2 diabetes (7). Recently, our study revealed that pharmacological inhibition of CRLs effectively reduced hyperglycemia in obese mice (8). We have shown that Cul3-containing RING E3 ligase (CRL3) mediates IRS protein turnover in hepatocytes and pan neddylation inhibitors (i.e. MLN4924, TAS4464) stabilize IRS protein to increase hepatic insulin sensitivity and reduce gluconeogenesis (8). In this study, we report that pan neddylation inhibitors also exhibit an insulinotropic effect, which is also mediated by CRL3 inhibition in β cells. We demonstrate that DI-1859, a selective Cul3 neddylation inhibitor (9), simultaneously promotes insulin secretion in β cells and IRS stabilization in liver and muscle cells. This dual action underlies the rapid and potent glucose lowering effect of DI-1859 in obese mice.

## Methods

### Cell culture experiments

AML12 cells, a gift from Dr. Yanqiao Zhang (Northeast Ohio Medical University, Rootstown, OH), were cultured in DMEM supplemented with 10% Fetal Bovine Serum (FBS) and insulin-transferrin-selenium (ITS, Cat. #: 41400-045, ThermoFisher Scientific, Waltham, MA). C2C12 myoblasts (American Type Culture Collection, Manassas, Virginia) were cultured in DMEM supplemented with 10% FBS and 1% penicillin–streptomycin. To induce differentiation into myotubes, C2C12 cells were switched to differentiation medium (DMEM supplemented with 2% horse serum) for 6 days (differentiation medium was replaced every two days). INS-1 832/13 cells were a gift from Dr. Christopher Newgard (Duke University, Durham, North Carolina) (10). INS-1 832/13 cells were cultured in RMPI1640 medium (12.5 mM glucose, supplemented with 2 mM L-glutamine, 1 mM sodium pyruvate, 10 mM HEPES, pH 7.4, 0.05 mM β-mercaptoethanol, penicillin–streptomycin, and 10% FBS). For insulin secretion, cells were cultured in poly-L-lysine-coated culture plates in RPMI1640 medium until full confluence. Cells were cultured in glucose and phenol red-free DMEM (Gibco, Cat. #: A14430-01, supplemented with 2.5 mM glucose, 2 mM L-glutamine, 1 mM sodium pyruvate, 10 mM HEPES, pH-7.4, 0.05 mM β-mercaptoethanol, 1% penicillin–streptomycin) overnight. The next morning, pre-treatments were performed and cells were washed 3 times with Krebs-Ringer Buffer (KRB) HEPES-buffered Solution (Cat. #: J67795.K2, ThermoFisher Scientific, supplemented to a final concentration of 3.5 mM glucose and 0.2% bovine serum albumin) and then cultured in the same KRB solution containing either 3.5 mM or 10 mM glucose with the same treatments. An aliquot of culture medium was collected for insulin measurement and normalized to total cellular protein.

### Mice and treatments

WT male C57BL/6J mice were purchased from the Jackson Laboratory (Bar Harbor, ME). Western diet (Cat. #: TD. 88137, Inotiv, Inc. Indianapolis, IN) contains 42% fat calories and 0.2% cholesterol. Mice were housed in micro-isolator cages with corn cob bedding under 7 am - 7 pm light cycle and 7 pm -7 am dark cycle. STZ was prepared in Na-Citrate solution (pH 4.5). Male C57BL/6J mice at 8 weeks old were i.p. injected with STZ at 50 mg/kg once daily for 5 consecutive days. After 2 weeks, hyperglycemia was confirmed and further experiments were performed. Euthanasia was performed with isoflurane inhalation. All animal protocols were approved by the Institutional Animal Care and Use Committee at the University of Oklahoma Health Sciences (protocol #22-072-EAFHI).

### Hyperinsulinemic euglycemic clamp

Hyperinsulinemic euglycemic clamps were performed by the Vanderbilt Mouse Metabolic Phenotyping Center (MMPC). Male DIO C57BL/6J mice (Strain #: 380050, 16 weeks old) were purchased from the Jackson Lab (Bar Harbor, ME) and shipped directly to the MMPC facility. Detailed procedure was described previously (11). All procedures required for the hyperinsulinemic euglycemic clamp were approved by the Vanderbilt University Animal Care and Use Committee.

### Human islet culture and insulin secretion

Freshly isolated human islets were purchased from Prodo Laboratories Inc. (Aliso Viejo, CA). Islets were cultured in Prodo islet complete standard culture medium (Cat. #: PIM-CS001GMP), supplemented with a glutamine/glutathione mixture, an antibiotics combination of Amphotericin B, Ciprofloxacin and Gentamicin, and heat inactivated human AB serum. The next day, islets were washed once with G55 3 mM medium. The G55 3 mM medium is made by first mixing 500 ml Ham’s F-10 nutrient mix medium (ThermoFisher Scientific, Cat. #: 11550043) and 500 ml no glucose DMEM (ThermoFisher Scientific, Cat. #: A1443001). Then, 600 mg sodium bicarbonate and 110 mg CaCl_2_.2H_2_O was added to the medium and pH was adjusted to 7.4. The G55 3 mM medium contains 3 mM glucose. The islets were then cultured in G55 3 mM medium in 24-well plates with Millipore Millicell 12 mm cell culture insert (Cat. #: PIXP01250, Sigma, St. Louis, MO) at ∼70 IEQ/well overnight. The next morning, insulin secretion assay was performed as described under INS-1 832/13 β cells.

### Glucose tolerance test (GTT) and mixed meal tolerance test (MMTT)

Mice were injected intraperitoneally 2 mg/kg glucose for GTT. Mice were orally administered 200 ml of Ensure Plus Nutrition Shake (Abbott Labs, USA; Per 8 oz, consists of 11 g total fat, 47 g total carbohydrates, 16 g total protein, and various vitamins plus minerals) for MMTT.

### Microscopy

For F-actin staining, fixed cells were incubated for 1 hour at room temperature with Phalloidin-California Red Conjugate (Santa Cruz Biotechnology, Inc. Cat. #: sc-499440, 1:300 dilution) in PBS. Coverslips were mounted using a DAPI-containing mounting medium (Cat. #: H-1200, Vector laboratories, Inc. Newark, CA). Images were acquired with a ThermoFisher EVOS M5000 fluorescence microscope.

### Western blotting

Protein lysates were prepared in 1XRIPA buffer containing 1% SDS and a cocktail of protease inhibitors and used for SDS-PAGE and immunoblotting.

### Measurement of mitochondria DNA (mtDNA) copy number

Total genomic DNA from INS-1 832/13 cells was purified by phenol-chloroform extraction followed by ethanol precipitation. The mtDNA copy number was measured with Absolute Rat Mitochondrial DNA Copy Number Quantification kit (Cat. #: R8948, ScienCell Research Laboratories, Inc. Carlsbad, CA).

### Co-immunoprecipitation and proteomics

INS-1 832/13 cells were cultured in 15 cm culture dish and infected with Ad-Cul3-FLAG or Ad-RhoA-FLAG (IP sample). The negative control (NC) was infected with Ad-Null. After 24 hours, both IP and NC lysates were subjected to anti-FLAG magnetic beads immunoprecipitation. The precipitated proteins were used for the protein-protein interaction profiling using the IP-works service provided by MSBioworks LLC (Ann Arbor, MI). Quantitative Proteomics was performed in the Multiplexing Protein Analysis Laboratory at Oklahoma Nathan Shock Center of Excellence in Aging Research. The mass spectrometry methods are described in Supplementary Methods.

### U-C_13_-glucose metabolic tracing through glycolysis

INS-1 832/13 cells were first cultured in DMEM (containing 3.5 mM glucose, 2 mM L-Glutamine, 1 mM sodium pyruvate, 10 mM HEPES, PH7.4.) overnight. The next morning, cells were treated with either vehicle (DMSO) or DI-1859 (100 nM) for 1 hour. Cells were then cultured in the same DMEM with added 10 mM U-C_13_-glucose for 5 hours with the same treatments. Cell pellets were collected and detection of glycolysis intermediate metabolites was performed at Creative Proteomics (Shirley, NY). Detection methods are described in Supplementary Methods.

### Seahorse flux assay of mitochondrial respiration and glycolysis analysis

Mitochondrial respiration and glycolysis in INS-1 832/13 cells were measured with Seahorse XFe96 Analyzer (Agilent Technologies, CA) following the manufacturer’s user guide for Mito Stress Test and Glycolysis Stress Test. All mitochondrial respiration rates were automatically calculated by the Seahorse XF Cell Mito Stress Test Report Generator and normalized to the cell number.

### Statistics

Unpaired or paired Student’s t-test was used for 2 group comparison. For multiple-group comparison, one-way or two-way ANOVA and Tukey post hoc test were used. A p < 0.05 was considered statistically significant.

**Note.** Please refer to Supplementary Methods for additional experimental details.

## Results

### Pan neddylation inhibitors exhibit dual insulin sensitization and insulinotropic effect

During hyperinsulinemic euglycemic clamp, acute MLN4924 treatment increased glucose infusion rate, reduced endogenous glucose production, and increased glucose uptake in skeletal muscle and white adipose in obese mice (**Fig 1A-E**). Unexpectedly, blood insulin concentration was significantly elevated by MLN4924 treatment (**Fig 1F**), which indicates a possible insulinotropic effect of MLN4924. While we could not clearly separate the possible insulin sensitization effect of MLN4924 in skeletal muscle in the presence of elevated blood insulin in vivo, we next studied the MLN4924 effect in culture differentiated C2C12 myotubes. Similar to the insulin sensitization effects of MLN4924 in hepatocytes (8), MLN4924 treatment stabilized IRS1 protein, enhanced AKT phosphorylation, and potentiated insulin-stimulated glucose uptake in C2C12 myotubes (**Fig 2A-C**). MLN4924 did not lower glucose in STZ-treated mice (**Fig 2D-E**), suggesting that the hypoglycemic effect of MLN4924 is largely insulin-dependent.

**Figure 1.**
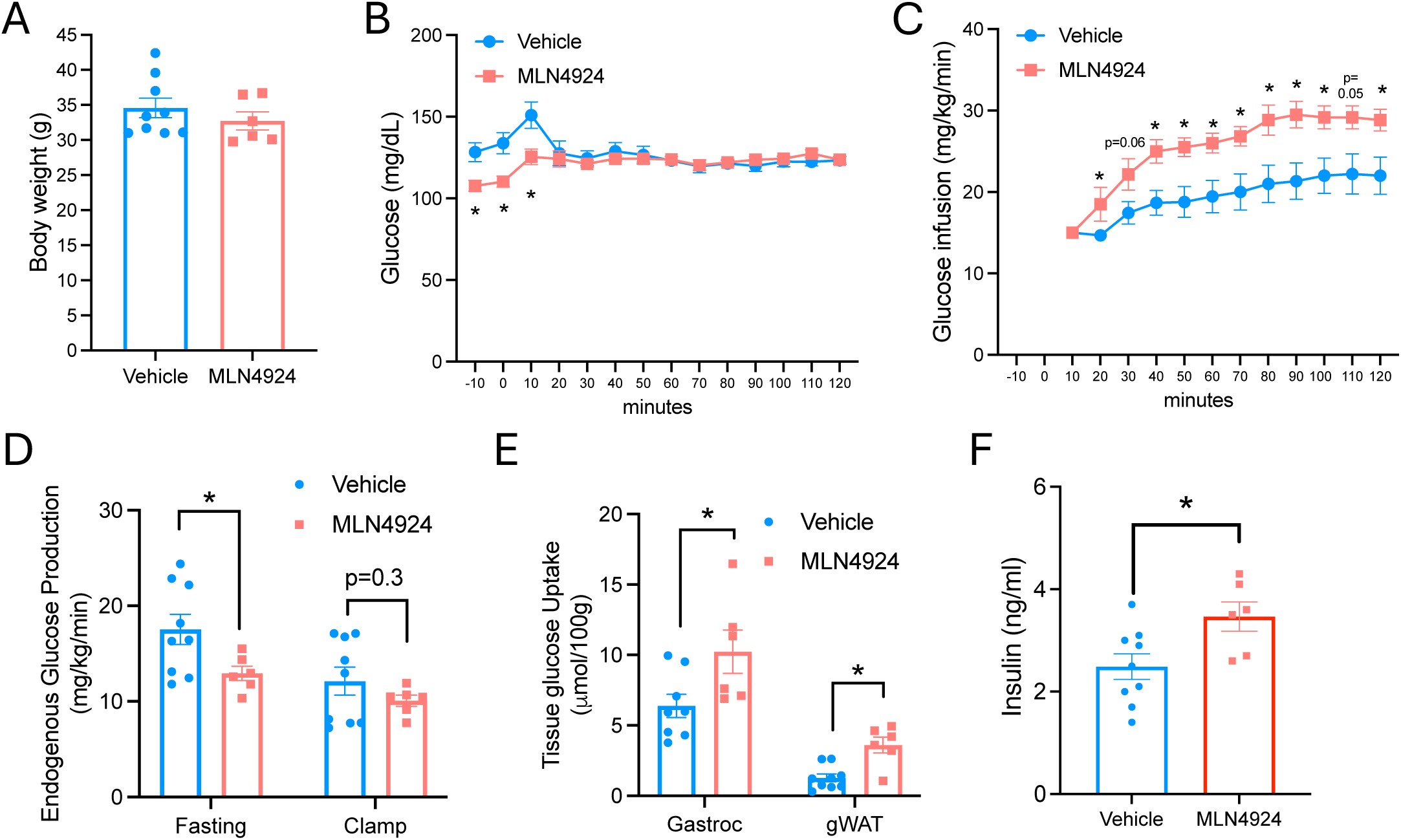
Hyperinsulinemic euglycemic clamp analysis of the effects of MLN4924 on whole body insulin action in diet-induced obese (DIO) mice. Male C57BL/6J DIO mice at 16 weeks of age were treated with i.p. injection of 60 mg/kg MLN4924 or vehicle (10% 2-hydroxypropyl-β-cyclodextrin) at 5 pm on day 1 and 9 am on day 2. Mice were fasted for ∼5 hours after the 2^nd^ treatment on day 2 and hyperinsulinemic euglycemic clamp was initiated. n=9 for the vehicle group and n=6 for the MLN4924 group. **A.** Body weight on the day of clamp. **B.** Blood glucose 10 mins before and during clamp. **C.** Glucose infusion rate. **D.** Endogenous glucose production. **E.** Tissue glucose uptake. Gastroc: gastrocnemius muscle. gWAT: gonadal white adipose tissue. **F.** Blood insulin at the end of clamp. All results are expressed as mean±SEM. Unpaired t-test was used for calculation of p values. A “*” indicates p< 0.05.

**Figure 2.**
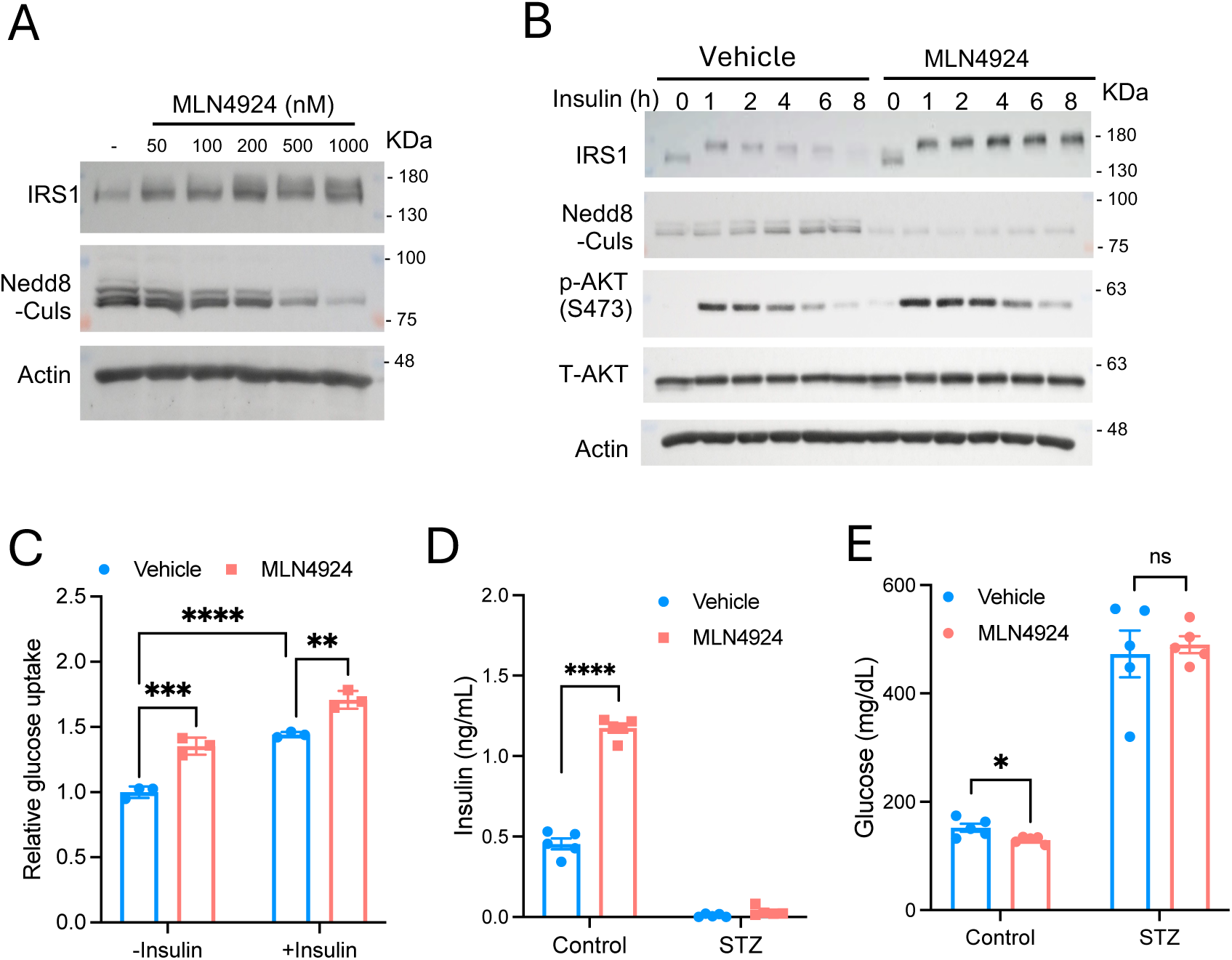
MLN4924 enhances insulin signaling and glucose uptake in muscle cells. **A.** Western blot. C2C12 myotubes were serum starved for 16 hours and treated with increasing concentrations of MLN4924 for 8 hours. **B.** Western blot. C2C12 myotubes were serum starved for 16 hours and then pre-treated with 500 nM MLN4924 or vehicle (DMSO) for 1 hour followed by 100 nM insulin stimulation in time course. **C.** C2C12 myotubes were serum starved for 16 hours. Cells were then pre-treated with 500 nM MLN4924 or vehicle (DMSO) for 6 hour followed by stimulation with 100 nM insulin for 1 hour as indicated. Glucose uptake was then performed with the Glucose Uptake-Glo assay kit. The result is from a representative experiment of 3 independent experiments with similar results. The data is expressed as the mean+SD of triplicates, with control (-insulin, vehicle) set as “1”. Two-way ANOVA and Tukey post hoc test was used for calculation of p values. **D, E.** Male STZ mice and non-STZ control mice were treated with vehicle or 60 mg/kg MLN4924 or vehicle (10% 2-hydroxypropyl-β-cyclodextrin) at 9 am and fasted for 6 hours. Blood samples were collected for insulin and glucose measurement. n=5. All results are expressed as mean±SEM. Unpaired t-test was used for calculation of p values. For all panels, “*”, <0.05; “**”, <0.01; “***”, <0.001; “****”, <0.0001. ns, not significant, p>0.05.

A single MLN4924 treatment in chow-fed mice lowered blood glucose and increased blood insulin and c-peptide (**Fig 3A-C**). Strong insulinotropic effect was also seen when chow-fed mice were treated with another pan neddylation inhibitor TAS4464 (**Fig 3D-E**). In an oral mixed meal tolerance test (MMTT), TAS4464 treatment increased blood insulin without potentiating postprandial glucagon-like peptide-1 (GLP-1) secretion (**Fig 3F-H**, **Supplementary Fig 1A-D**). In INS-1 832/13 β cells (10), MLN4924 treatment rapidly repressed Cullin neddylation and potentiated glucose-stimulated insulin secretion (**Fig 3I-J**), suggesting a direct insulinotropic effect of MLN4924 in β cells.

**Figure 3.**
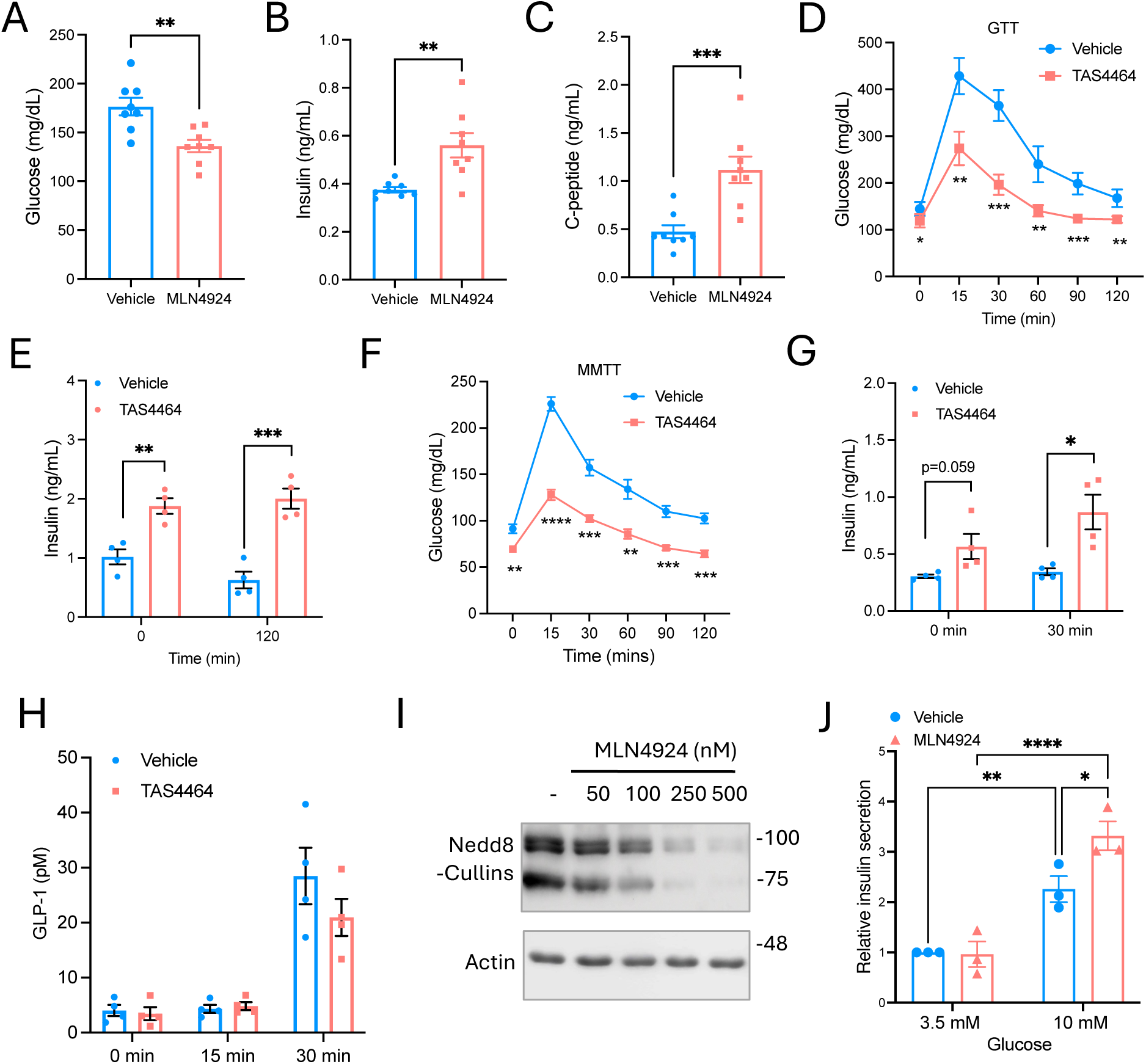
Neddylation inhibitors promote insulin secretion in mice and cultured INS-1 832/13 β cells. **A-C.** Male C57BL/6J mice at ∼10 weeks of age were i.p. injected with vehicle or 60 mg/kg MLN4924 or vehicle (10% 2-hydroxypropyl-β-cyclodextrin) at 9 am and then fasted for 6 hours. Blood glucose, insulin and c-peptide were measured. n=8. **D-E.** Glucose tolerance test (GTT). Male C57BL/6J mice at ∼10 weeks of age were i.p. injected with vehicle or 45 mg/kg TAS4464 or vehicle (10% 2-hydroxypropyl-β-cyclodextrin) at 9 am and then fasted for 6 hours. GTT was then performed. Blood samples were collected at 0 and 120 minutes for insulin measurement. n=4. **F-H.** Mixed meal tolerance test (MMTT). Male C57BL/6J mice at ∼10 weeks of age were fasted overnight, and i.p. injected with vehicle or 45 mg/kg TAS4464 or vehicle (10% 2-hydroxypropyl-β-cyclodextrin) at 9 am and MMTT was initiated at 11 am. Blood samples were collected for insulin and GLP-1 measurement. n=4. **I.** Western blot. INS-1 832/13 cells were treated with MLN4924 in dose response for 30 minutes. Anti-Nedd8 antibody was used to detect neddylated Cul proteins. **J.** Glucose-stimulated insulin secretion. INS-1 832/13 cells were cultured in 2.5 mM glucose DMEM overnight and then pre-treated with vehicle (DMSO) or 100 nM MLN4924 for 1 hour the next morning. Cells were then cultured in KRB buffer containing 3.5 mM or 10 mM glucose for insulin secretion assay. The results are the mean of 3 independent experiments with the vehicle+3.5 mM glucose group set as “1”. All results are expressed as mean±SEM. The p values were calculated using unpaired t-test for A-H, and 2-way ANOVA and Tukey post hoc test for J.

### Selective Cul3 neddylation inhibitor DI-1859 promotes insulin sensitization and insulin secretion and lowers blood glucose in obese mice

Pan neddylation inhibitors MLN4924 and TAS4464 potently inhibit the neddylation of all Cullin proteins (3). Recently, covalent binding of DCN1 in the neddylation E2 complex by DI-1859 was discovered to selectively inhibit Cul3 neddylation (9). Our prior study identified that Cul3 mediated IRS protein turnover and the insulin sensitization effect of pan neddylation inhibitors in hepatocytes (8), which prompted us to investigate if DI-1859 possessed glucose lowering effect. Interestingly, a single DI-1859 administration in obese mice significantly decreased blood glucose (**Fig 4A-B**), elevated blood insulin and c-peptide (**Fig 4C-D**), and improved glucose tolerance (**Fig 4E**). DI-1859 treatment did not lower blood glucose in STZ-treated mice (**Fig 5A-B**), suggesting its effect is largely insulin-dependent. DI-1859 selectively inhibited Cul3 neddylation and stabilized IRS1 protein in liver AML12 cells (**Fig 5C-D**). Similarly, DI-1859 also inhibited Cul3 neddylation and stabilized IRS1 protein in C2C12 myotubes (**Fig 5E**). In both cell types, DI-1859 treatment enhanced/prolonged insulin stimulation of AKT phosphorylation (**Fig 5D-E**). These results are consistent with the reported role of Cul3 in promoting IRS protein stability and cellular insulin sensitivity (8). In summary, selective inhibition of Cul3 fully recapitulated the insulin sensitization and insulinotropic effects of pan neddylation inhibitors.

**Figure 4.**
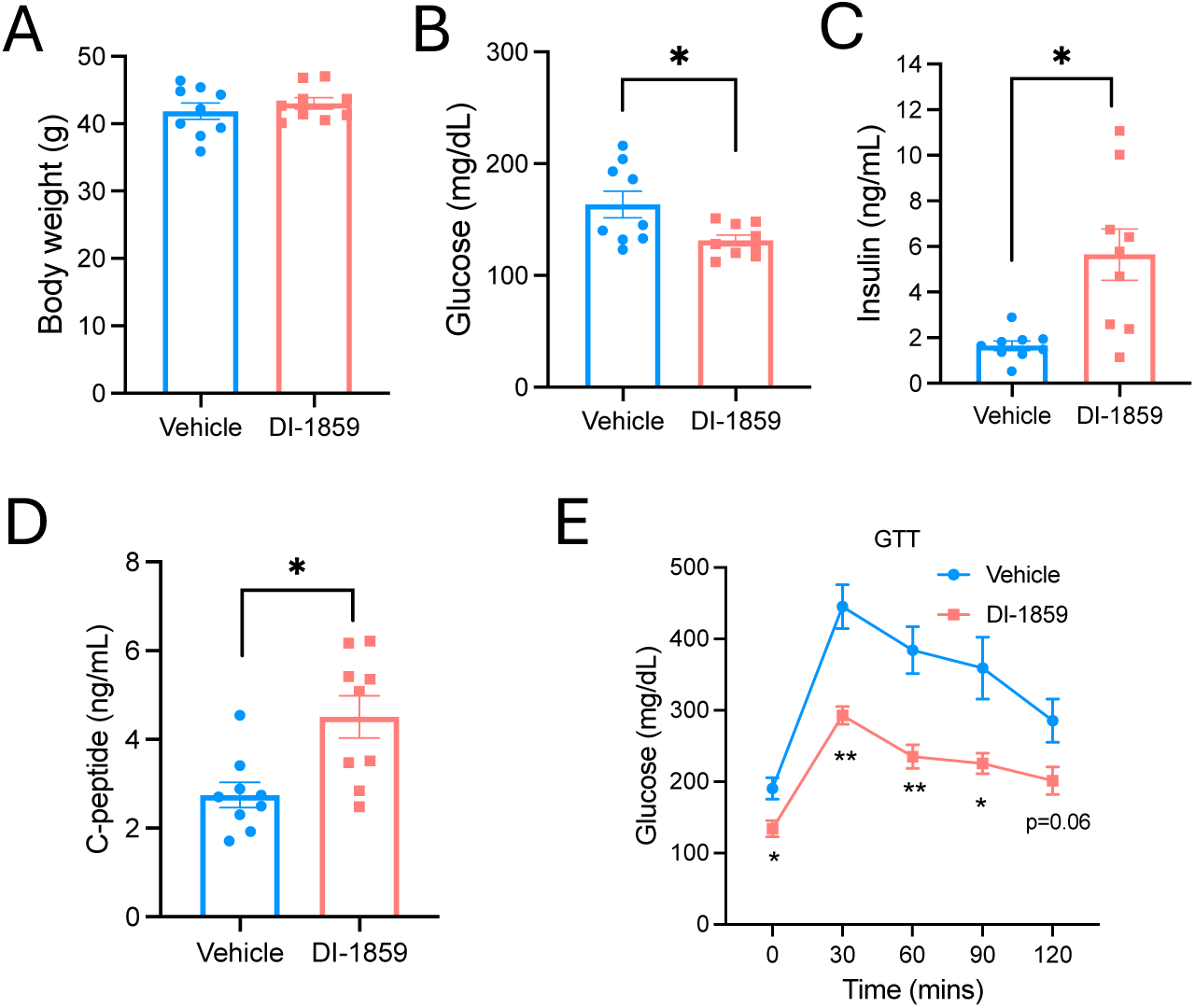
Selective Cul3 inhibitor promotes insulin secretion and lowers glucose in obese mice. Western diet-fed male C57BL/6J obese mice were i.p. injected with vehicle or 10 mg/kg DI-1859 or vehicle (10% 2-hydroxypropyl-β-cyclodextrin) at 9 am and blood samples were collected at 3 pm after a 6-hour fast. **A.** Body weight. **B.** Blood glucose. **C.** Blood insulin. **D.** Blood c-peptide. **E.** GTT in Western diet-fed male C57BL/6J obese mice treated with vehicle or DI-1859 for 6 hours. n=9 for A-D. n=4 for E. All results are expressed as mean ± SEM. Unpaired t-test was used for calculation of p values. For all panels, “*”, <0.05; “**”, <0.01.

**Figure 5.**
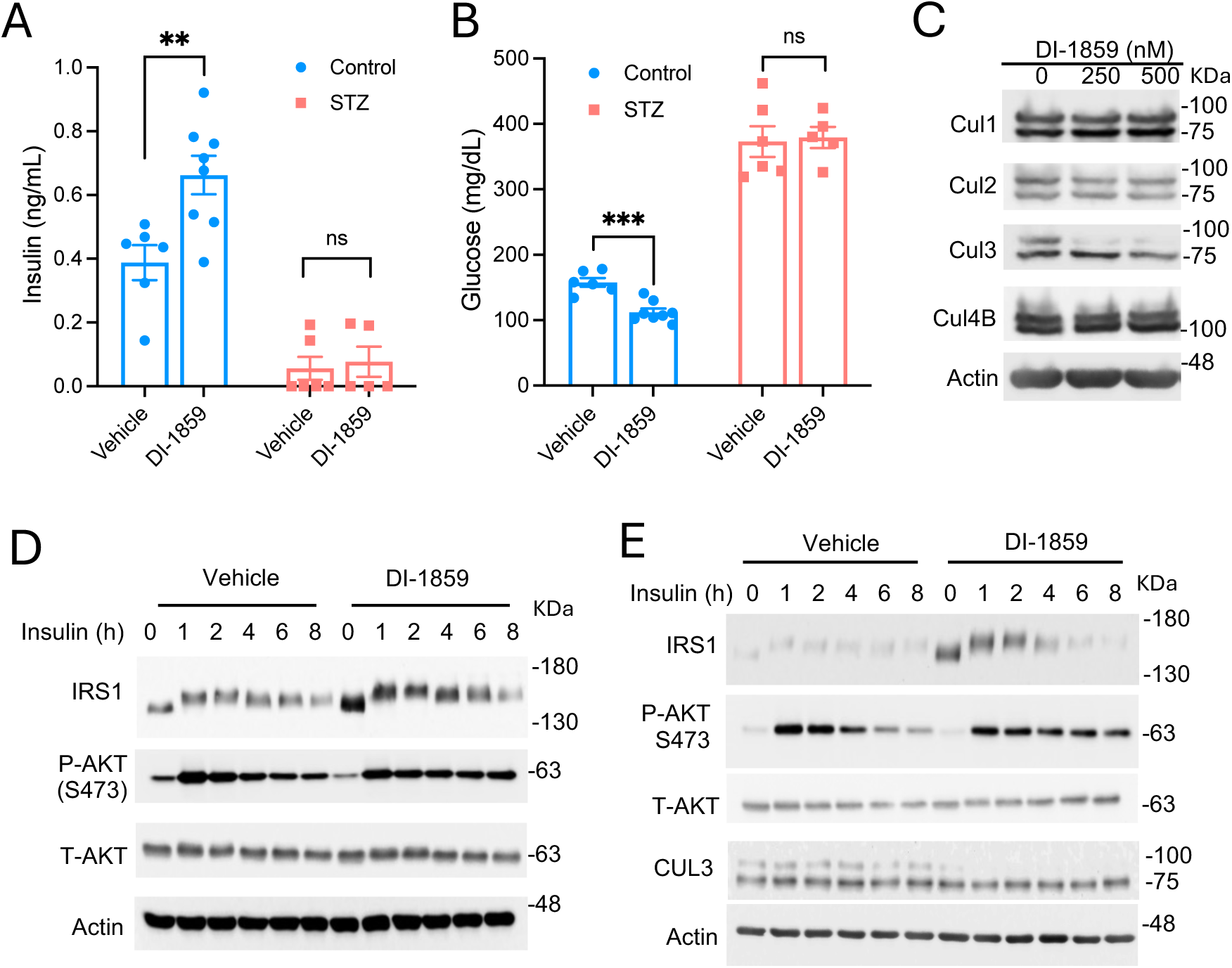
The glucose lowering effect of DI-1859 is dependent on insulin signaling axis. **A-B.** Male STZ mice and non-STZ control mice were treated with vehicle (10% 2-hydroxypropyl-β-cyclodextrin) or 10 mg/kg DI-1859 at 9 am and fasted for 6 hours. Blood samples were collected for insulin and glucose measurement. n=5-8. **C.** Western blot. AML12 cells were treated with DI-1859 for 2 hours. **D.** Western blot. AML12 cells were serum and ITS starved overnight. The next morning, all cells were pre-treated with 100 μg/ml cycloheximide and either vehicle (DMSO) or 250 nM DI-1859 for 1 hour and then stimulated with 100 nM insulin in time course. **E.** Western blot. C2C12 myotubes were serum starved overnight. The next morning, all cells were pre-treated with 100 μg/ml cycloheximide and either vehicle (DMSO) or 250 nM DI-1859 for 1 hour and then stimulated with 100 nM insulin in time course. A representative Western blot of at least 3 independent experiments is shown.

### Selective Cul3 inhibition promotes insulin secretion in cultured INS-1 832/13 β cells and human islets

DI-1859 treatment caused rapid and selective inhibition of Cul3 neddylation and potentiated glucose-stimulated insulin secretion in INS-1 832/13 β cells (**Fig 6A-B**). This was unlikely an off-target effect because knockdown of Cul3 also potentiated glucose-stimulated insulin secretion (**Fig 6C-D**). We next investigated the effect of DI-1859 in cultured human islets. Because the absolute insulin secretion varied significantly among different islet preparations, we plotted the result separately with the respective control set as “1”. We confirmed that DI-1859 treatment repressed Cul3 neddylation in human islets (**Fig 6E**). Exposure to 10 mM glucose increased insulin secretion by ∼ 3-fold over basal insulin secretion under 3.5 mM glucose condition (**Fig 6F**). DI-1859 treatment increased insulin secretion by ∼ 50% over vehicle-treated controls under both 3.5 mM and 10 mM glucose condition (**Fig 6G-H**). These results show that selective inhibition of Cul3 promotes insulin secretion in β cells.

**Figure 6.**
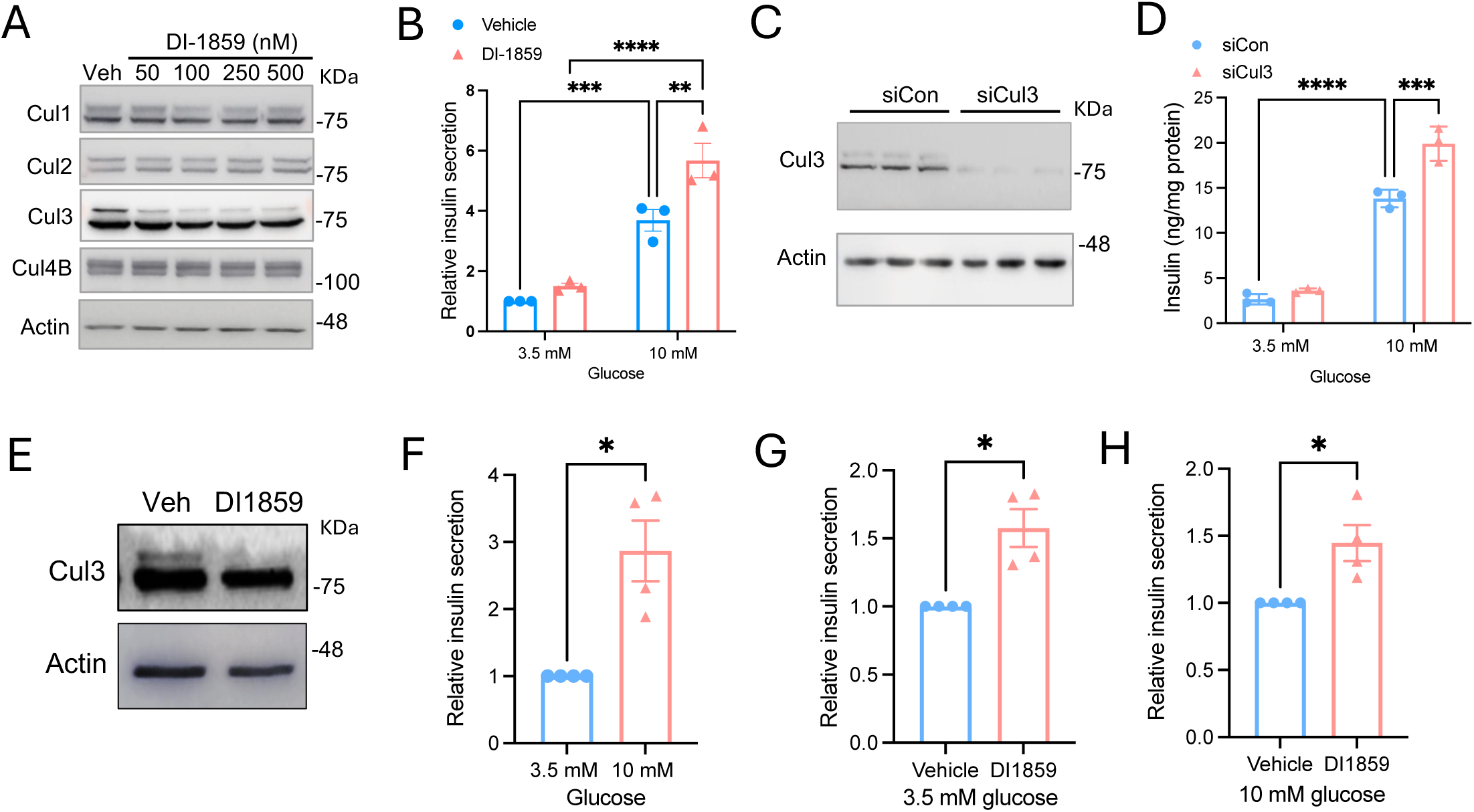
Selective Cul3 inhibition potentiates glucose-stimulated insulin secretion in INS-1 832/13 β cells and human islets. **A.** Western blot. INS-1 832/13 cells were treated with DI-1859 for 30 minutes. **B.** Glucose-stimulated insulin secretion. INS-1 832/13 cells were cultured in 2.5 mM glucose DMEM overnight and then pre-treated with vehicle (DMSO) or 100 nM DI-1859 for 1 hour the next morning. Cells were then cultured in KRB buffer containing 3.5 mM or 10 mM glucose for insulin secretion assay. The results are the mean±SEM of 3 independent experiments with the vehicle+3.5 mM glucose group set as “1”. **C.** Western blot. siRNA knockdown of Cul3 after 24 hours of siRNA transfection in INS-1 832/13 cells. Transfection was performed in triplicates. **D.** INS-1 832/13 cells were transfected with siRNA for 24 hours and then subject to glucose-stimulated insulin secretion assay. The results are from a representative experiment of 3 independent experiments with similar results. The assay was performed in triplicates and expressed as mean ±SD. The p values were calculated with 2-way ANOVA and Tukey post hoc test. **E-H.** DI-1859 effect on Cul3 neddylation (E) and insulin secretion (F-H) in human islets. Results are expressed as mean ±SEM with controls arbitrarily set as “1”. The p values were calculated with paired t-test. “*”, <0.05, “**”; <0.01; “***”, <0.001; “****”, <0.0001. ns. not significant, p>0.05.

### DI-1859 does not promote glycolysis or cellular bioenergetic activity

Because Cul3 inhibition may enrich its protein substrates, we next perform unbiased quantitative proteomics hoping to identify CRL3-targeted factors linked to the insulin secretion processes. This assay detected a total of 3952 proteins, among which 592 proteins were significantly upregulated (fold change cutoff: 1.2; p value: <0.05), while only 17 proteins were significantly downregulated (fold change cutoff: -0.8; p value <0.05) (**Supplementary Fig 2A**). Ingenuity Pathway Analysis (IPA) identified that several of the top upregulated pathways by DI-1859 treatment were related to cellular glucose metabolism (**Supplementary Fig 2B**), which was partly due to modestly increased glycolytic enzymes GPI, ALDOA, TPI1, GAPDH, PGK and PGAM1 (**Supplementary Fig 2C**). Other core glycolytic enzymes, including the rate-limiting enzyme PFKM, were not significantly altered by DI-1859 treatment (**Supplementary Fig 2C**). Given the critical role of glucose metabolism in insulin secretion (12), we next investigated if DI-1859-treated cells show increased glycolysis flux using U-C^13^-glucose tracing (**Fig 7A**). We found that M+0 and M+6 upper glycolytic intermediates (glucose, G6P, F6P) were similar between the two groups, suggesting equal entry of glucose into the upper glycolysis pathway (**Fig 7B**). The downstream M+3 3PG, pyruvate and lactate, which are considered key indicators of glycolysis flux, were not altered by DI-1859 treatment (**Fig 7B**). The only change we observed was a decrease of M+0 and M+6 FBP in DI-1859-treated cells. A decrease of these intermediates could indicate reduced glycolytic flux at the PFK step, which, however, was not supported due to the lack of accumulation of upper glycolysis intermediates (M+6 F6P and G6P) and no increased flux into the pentose phosphate pathway (PPP) (**Fig 7B-D**). Alternatively, the reduction of these intermediates could reflect a “pull” on FBP due to an increase of ALDOA without a significant impact on the downstream flux. More in line with the latter scenario, Seahorse flux analysis did not show altered ECAR in DI-1859-treated INS-1 832/13 β cells (**Fig 7E-F**), which provides functional evidence that DI-1859 does not promote overall glycolysis flux. We further evaluated mitochondrial bioenergetic function using Seahorse flux analysis and found no difference between the two groups (**Supplementary Fig 3A**). In comparison to the clear colocalization with mitochondrial protein TOM20 (positive control) of an engineered mitochondria-targeted protein mitoSTH (serves as a positive control), Cul3 showed a primarily cytosolic localization with no apparent mitochondrial localization (**Supplementary Fig 3B**). Furthermore, DI-1859 treatment did not have a significant effect on gross mitochondrial morphology or mitochondrial abundance (**Supplementary Fig 3C-D**). Taken together, we found no evidence that DI-1859 treatment increased glycolysis or cellular bioenergetic activity. Therefore, further validation of the elevated glycolytic enzymes as CRL3 substrates was not pursued in this study.

**Figure 7.**
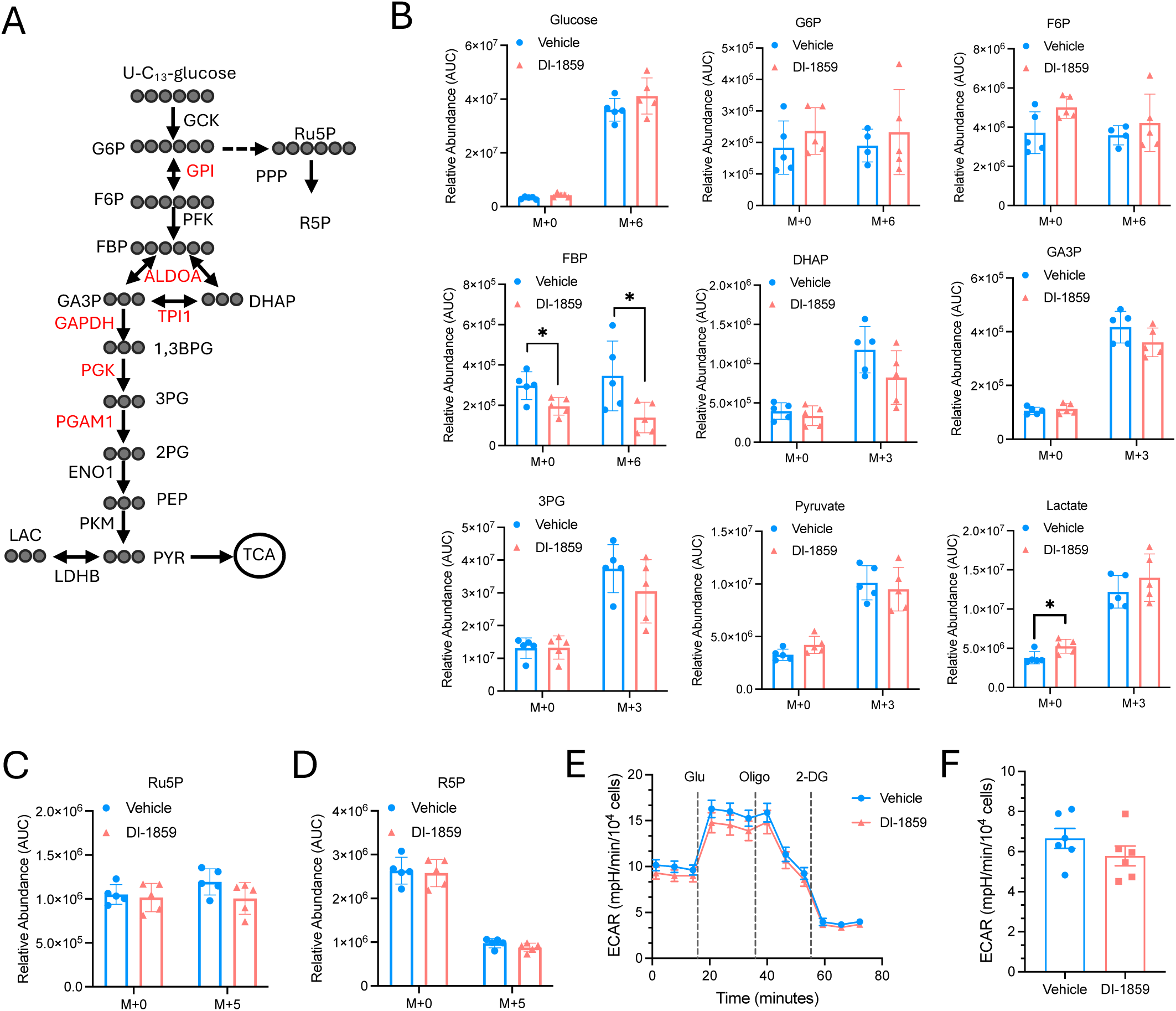
DI-1859 does not promote glycolysis INS-1 832/13 cells. **A.** Glycolysis pathway and pentose phosphate pathway (PPP). U-C_13_-glucose labeling of intermediate metabolites is illustrated. Red font indicates upregulated enzymes by DI-1859. **B-D.** Unlabeled (M+0) and C_13_-labeled metabolites by U-C_13_-glucose in metabolic flux tracing analysis. n=5. **E-F.** INS-1 832/13 cells. Glycolysis assay was performed by using Seahorse Flux analyzer. On the day of measurements, cells were pre-treated with Vehicle or 100 nM DI-1859 for 2 h in DMEM-XF (2.5 mM Glucose). Cells were then switched to Seahorse bicarbonate-free RPMI-XF (Aglient 103576-100, supplemented with 12.5 mM glucose, 2 mM L-glutamine, 1 mM sodium pyruvate, pH 7.4), with Vehicle or 100 nM DI-1859 treatment, and equilibrated at 37 °C non-CO2 incubator for 1 h. ECAR was monitored with injection of glucose (12.5 mM), oligomycin (1.5 μM), and 2-DG (50 μM). Quantitation of glucose stimulated ECAR is shown in “F”. n=6. All replicates are from independent wells of a culture plate. All results are expressed as mean ± SEM. The p values were calculated with t-test. “*”, <0.05.

### The CRL3-RhoA dependent regulation of cytoskeleton is involved in mediating the insulinotropic effect of DI-1859

We next performed a Cul3 co-IP proteomics to identify Cul3 interacting proteins that may be involved in regulation of insulin secretion (**Supplementary Fig 4A**). We found that 43 of the top 100 abundantly co-precipitated proteins were Cul3, the mammalian Cullin RING E3 ligase Rbx1, BTB domain-containing proteins and Kelch-like proteins that were predicted to serve as substrate receptors for Cul3 (2, 5), and all core subunits of the COP9 signalosome complex that mediates cullin de-neddylation (**Supplementary Table 1**). These data indicate that Cul3 protein is in relatively tight or constant association with its obligated substrate receptors and RING E3 ligase to form the core CRL3 complex. Its strong association with COP9 signalosome complex subunits may explain the rapid Cul3 de-neddylation upon DI-1859 treatment. Most other co-precipitated proteins, some of which are potential CRL3 substrates, are in relatively low abundance, possibly due to the transient nature of the CRL3-substrate interaction, which also implied that the Cul3 Co-IP likely missed a significant number of CRL3 substrates. Cross examination of the Cul3 Co-IP proteomics and the unbiased quantitative proteomics datasets identified 62 Cul3-associated proteins that were also significantly enriched in cells treated with DI-1859. The majority of these proteins are not core CRL3 components and could represent a list of potential CRL3 substrates in β cells. However, they did not offer apparent mechanistic clue to potentiated insulin secretion upon Cul3 inhibition (**Supplementary Fig 4B, Supplementary Table 2**).

Interestingly, we found that the obligated RhoA substrate receptors BACURD1, BACRUD2 and BACURD3 (BTB/POZ domain-containing adapter for CUL3-mediated RhoA degradation protein 1, 2 and 3) were abundantly co-precipitated with Cul3 (**Supplementary Table 1**), which is consistent with RhoA being a validated CRL3 substrate (13, 14). RhoA is a small GTPase that plays a key role in regulating actin-microtubule cytoskeleton formation and dynamic remodeling that controls cell shape and intracellular transport (15, 16). Studies have recognized that the dynamic cytoskeleton remodeling critically regulates basal and stimulated insulin secretion in β cells (17–20). Consistently, RhoA was among the top 50 of all 594 enriched proteins detected by quantitative proteomics in DI-1859 treated cells (**Fig 8A**). IPA analysis of the quantitative proteomics dataset showed that many upregulated pathways by DI-1859 were related to Rho GTPase signaling and cytoskeleton regulation (**Supplementary Table 3**).

**Figure 8.**
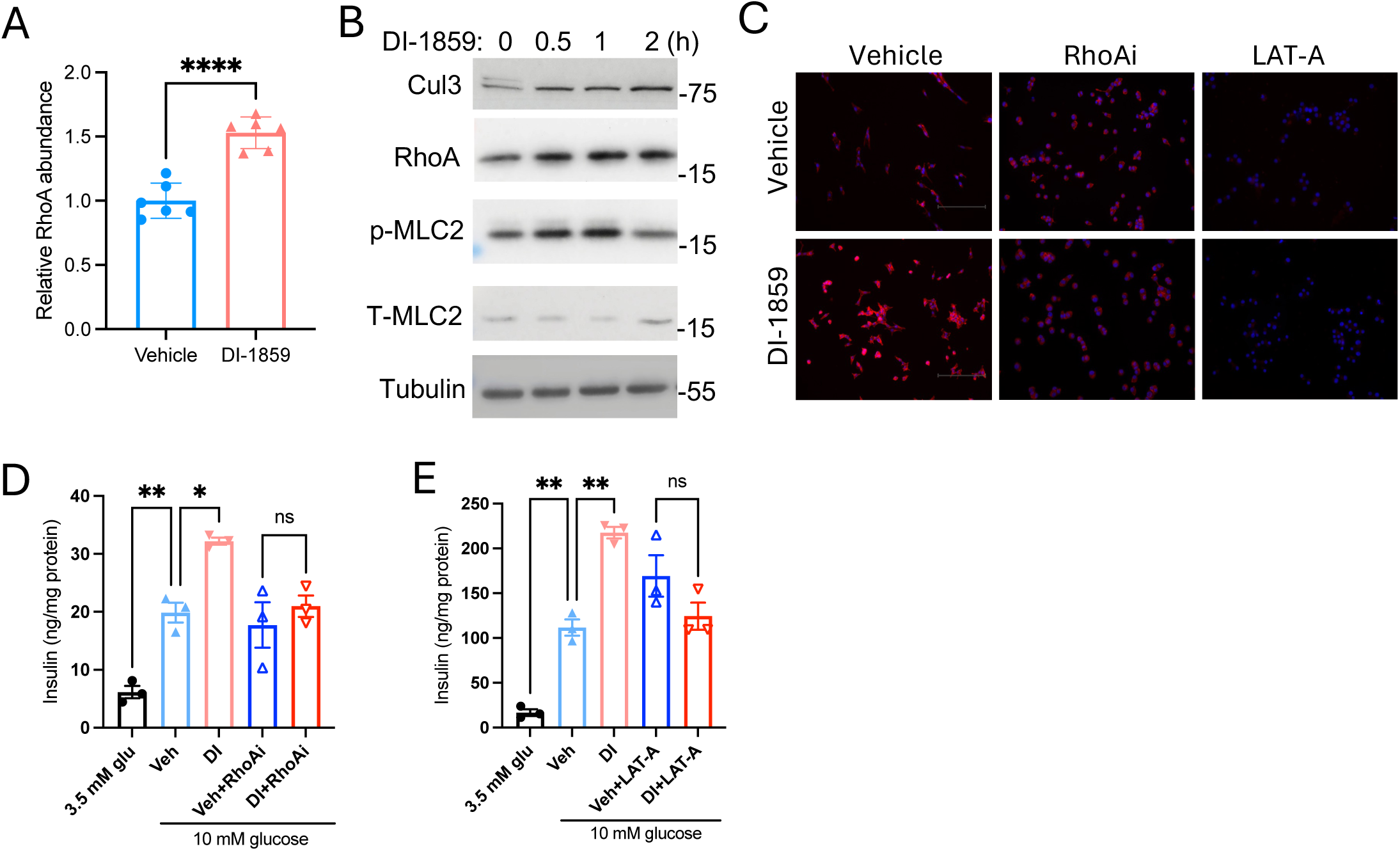
DI-1859 potentiation of insulin secretion involves activation of RhoA and cytoskeleton remodeling in INS-1 832/13 cells. **A.** RhoA protein detected by quantitative proteomics in INS-1 832/13 cells treated with DI-1859 for 3 hours. **B.** Western blot. INS-1 832/13 cells were pre-treated with treated with DI-1859 (100 nM). **C.** Immunofluorescent staining of F-actin by phalloidin (red) in INS-1 832/13 cells. Cells were pre-treated with 100 ng/ml of RhoA inhibitor (C3, RhoAi) or 0.5 μM of Latrunculin A (LAT-A) as indicated for 1 hour followed by treatment with DI-1859 (100 nM) or vehicle for 1 hour. Nuclei were stained with DAPI (blue). Scale bar=125 μm. **D-E.** INS-1 832/13 cells were cultured in 2.5 mM glucose DMEM overnight and then pre-treated with vehicle (DMSO, Veh) or 100 nM DI-1859 (DI) and 100 ng/ml of RhoA inhibitor (C3, RhoAi) or 0.5 μM of Latrunculin A (LAT-A) as indicated for 1 hour the next morning. Cells were then cultured in KRB buffer containing 3.5 mM or 10 mM glucose under the same treatment condition as indicated for insulin secretion assay. The results are from a representative experiment of 3 independent experiments with similar results. The assay was performed in triplicates. All results are expressed as mean ± SEM. The p values were calculated with t-test for A and one-way ANOVA and Tukey post hoc test for D-E. “*”, <0.05, “**”, <0.01; “***”, <0.001; “****”, <0.0001. ns. not significant.

Consistently, DI-1859 treatment increased RhoA and the phosphorylation of RhoA target myosin light chain (MLC), a key activator of F-actin cytoskeleton remodeling, and the staining intensity of F-actin in INS-1 832/13 β cells (**Fig 8B-C**).

Treating INS1 832/13 β cells with a RhoA inhibitor (C3), which prevented DI-1859 induction of F-actin formation (**Fig 8C**), fully abolished the insulinotropic effect of DI-1859 (**Fig 8D**). Based on the knowledge that RhoA elicits a cellular effect via close association with its downstream effector proteins, we next preformed RhoA Co-IP proteomics. We found that RhoA was abundantly associated with guanine nucleotide exchange factors (GEF) Rap1gds1 and Arhgef1 (RhoA activators) and Rho GDP dissociation inhibitor (GDI) Arhgdia (RhoA inhibitor), which are known to mediate the dynamic cycling of RhoA between the active and inactive form (**Supplementary Table 4**). Interestingly, γ-Actin, a key cytoskeleton component, was also abundantly co-precipitated with RhoA (**Supplementary Table 4**). This is in line with prior knowledge that many GEFs and GDIs bind cytoskeleton and bridge RhoA and the cytoskeleton (21). Interestingly, key components and regulators of the actin-microtubule cytoskeleton also abundantly co-precipitated with Cul3 (**Supplementary Table 5**). These data suggest that RhoA and its regulators and effectors may be in proximity to allow for rapid and localized control of dynamic cytoskeleton remodeling. Based on these findings, we further treated INS-1 832/13 β cells with Latrunculin A (LAT-A) that directly disrupts F-actin cytoskeleton assembly (**Fig 8C**). Indeed, LAT-A fully abolished the insulinotropic effect of DI-1859 (**Fig 8E**). These findings together suggest that the insulinotropic effect of DI-1859 involves CRL3-RhoA regulation of cytoskeleton.

## Discussion

In this study we show that pan-neddylation inhibitors simultaneously promote insulin sensitization and insulin secretion, which helps explain their potent and rapid glucose lowering effect in obese mice. In our previous study, Cul3, but not other Cullin proteins, was shown to regulate IRS protein stability in hepatocytes (8). In this study, we further showed that the Cul3-mediated regulation of IRS stability was conserved in muscle cells, and selective inhibition of Cul3 by DI-1859 enhanced insulin signaling in both liver cells and myocytes. Intriguingly, we found that the insulinotropic effect of neddylation inhibitors was also mediated by Cul3 in β cells, which is the molecular basis underlying the dual action of DI-1859. Increased insulin secretion likely contributes significantly to the overall glucose reduction, although we cannot clearly separate the insulin sensitization and insulinotropic effect of DI-1859 in vivo. To date, the therapeutic potential of targeting Cullin neddylation in various pathological settings has been mostly investigated using pan neddylation inhibitors. Determining if certain therapeutic benefits of pan neddylation inhibitors are mediated by a specific Cullin protein not only improves the mechanistic understanding but also establishes the molecular basis for developing selective Cullin targeting strategies. In this study, we provide proof-of-principle evidence that selective inhibition of Cul3 neddylation is a feasible and effective strategy to attenuate hyperglycemia.

Our mechanistic study suggests that the insulinotropic effect of Cul3 inhibition by DI-1859 involves RhoA-mediated cytoskeleton regulation, supported by the finding that both RhoA inhibitor and cytoskeleton inhibitor fully prevent the insulinotropic effect of DI-1859 under high glucose exposure in β cells. The critical role of RhoA regulation of cytoskeleton has been well documented in various cell types (21). Consistent with prior knowledge, our Co-IP proteomics showed that RhoA was abundantly associated with GEF and GDI (21, 22). In addition, RhoA undergoes prenylation, a posttranslational modification that anchors RhoA to the plasma membrane where locally activated RhoA interacts with its regulators and effectors that bridge RhoA and the cytoskeleton. The role of RhoA in modulating exocytosis is complex and often found to be context dependent. It has been reported that the formation of dense F-actin polymers, a process that can be promoted by high sustained RhoA activity, can serve as a physical barrier for reserved insulin granules to gain access to the cell periphery, therefore limiting insulin secretion under basal non-stimulated conditions (19, 23). Paradoxically, high glucose exposure has also been associated with increased F-actin formation, and F-actin depolymerization impairs glucose-stimulated insulin secretion (24, 25). A unified model suggests that glucose-stimulated insulin secretion requires localized and concerted Actin polymerization and depolymerization, and this spatial and temporal actin remodeling controls the transport of insulin granules to the plasma membrane and exocytosis of insulin (26). In this study, we found that a short-term treatment of RhoA inhibitor and cytoskeleton assembly inhibitor LAT-A did not affect the magnitude of high glucose-stimulated insulin secretion but fully abolished the potentiating effect of DI-1859, suggesting the potentiating effect of DI-1859 in the presence of high glucose depends on both RhoA activation and undisrupted actin cytoskeleton that is amenable to dynamic remodeling. In our mechanistic study, we mainly utilized validated pharmacological RhoA inhibitor, which allowed rapid RhoA inhibition for study of glucose-stimulated insulin secretion in a short period of time. We found that constitutive knockdown of RhoA using siRNA for extended period (24-48 hours) significantly decreased cell growth and altered cell morphology, which was deemed unsuitable for further study of the role of RhoA in insulin secretion.

Future study is still needed to obtain molecular details on how CRL3-RhoA axis may control spatial and temporal cytoskeleton remodeling and the differential influence of such changes on basal and stimulated insulin secretion. Interestingly, Cul3 Co-IP proteomics analysis reveals that Cul3 is associated with many actin-microtubule cytoskeleton components, implying a significant cytoskeleton localization of the cytosolic pool of CRL3. This may place CRL3 in proximity with cytoskeleton-associated RhoA. While studies have reported Cul3 regulation of cytoskeleton remodeling via RhoA-dependent regulation (13, 27), direct CRL3 regulation of the turnover of cytoskeleton components has been reported (28). This study has identified a list of Cul3-associated substrate receptors and potential CRL3 substrates in β cells. Future study to characterize the substrate profile of these CRL3 obligatory receptors may add new mechanistic insights into the roles of CRL3 in regulation of insulin secretion and other aspects of β cell biology.

## Author Contributions

LG, LX, MNH, YD, TW, and TL performed the experiments and data analysis. TL supervised the study and wrote the manuscript. Timothy Yuze Wu is a summer student from Weston High School, Weston, MA.

## Guarantor Statement

T.L. is the guarantor of this work and, as such, had full access to all the data in the study and takes responsibility for the integrity of the data and the accuracy of the data analysis.

