## Supplementary Figures for "A selective Cullin 3 RING E3 ligase inhibitor attenuates hyperglycemia via dual insulin sensitizing and insulinotropic action"

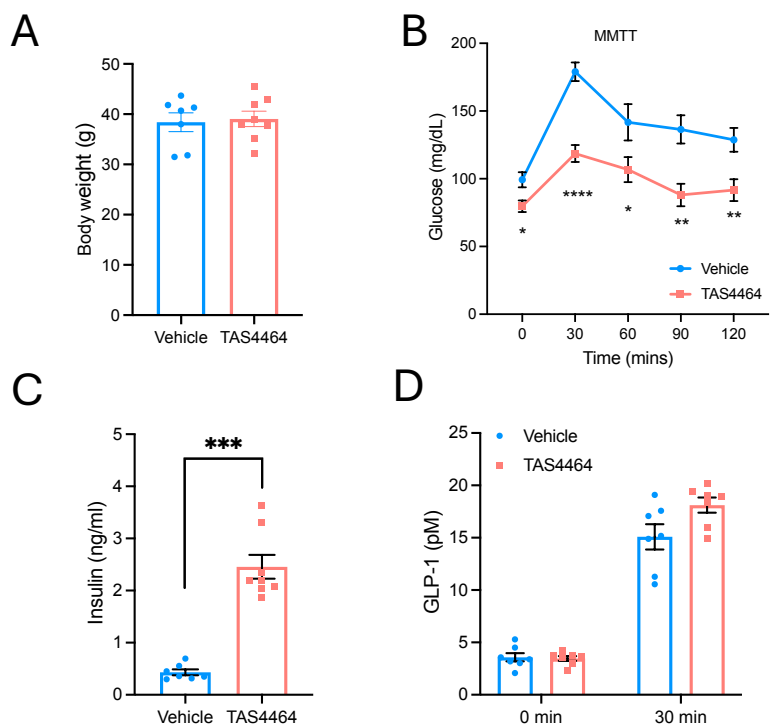

**Supplementary Figure 1. TAS4464 stimulates insulin secretion independent of GLP-1 in obese mice.** Male C57BL/6J mice at 8 weeks of age were fed WD for 8 weeks. These mice were fasted overnight, and i.p. injected with vehicle or 45 mg/kg TAS4464 at 9 am and mixed meal tolerance test (MMTT) was initiated at 2 pm on the same day. n=7-8. **A.** Body weight at the time of MMTT. **B.** Blood glucose. **C.** Blood insulin at 0 min. **D.** Blood GLP-1 at 0 min and 30 min during MMTT. Results are expressed as mean ± SEM. Unpaired t-test was used for calculation of p values. For all panels, “\*”, <0.05; “\*\*”, <0.01; “\*\*\*”, <0.001; “\*\*\*\*”, <0.0001.

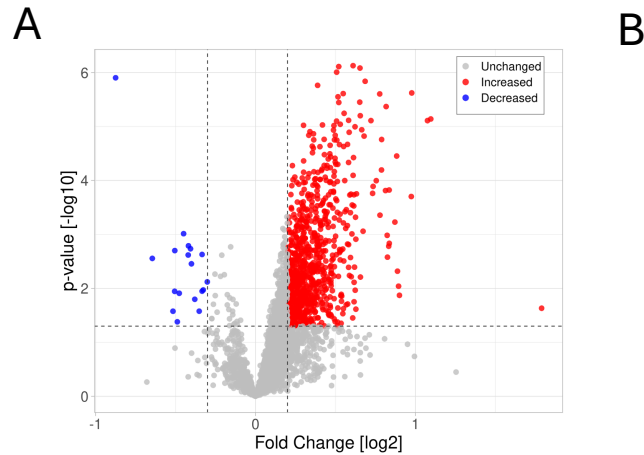

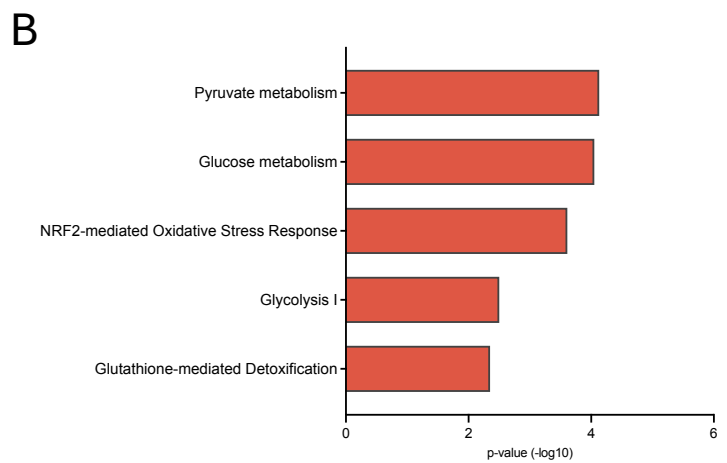

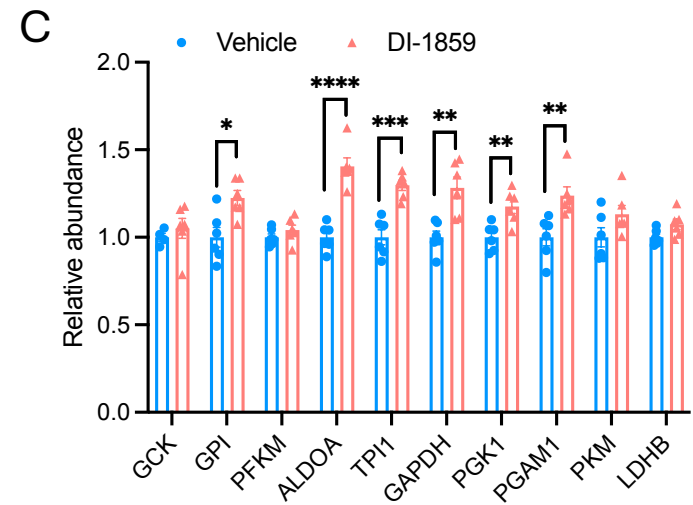

**Supplementary Figure 2. DI-1859 treatment enriches proteins involved in glucose metabolism in INS-1 832/13 $\beta$ cells.** INS-1 832/13 cells were cultured in DMEM containing 2.5 mM glucose overnight and treated with vehicle or 100 nM DI-1859 for 3 hours. Samples (6 replicates per condition) were collected for proteomics analysis. **A.** Volcano plot showing increased and decreased proteins by DI-1859 treatment. **B.** Ingenuity pathway analysis. Significantly upregulated pathways related to glucose metabolism and NRF2 activation by DI-1859. Increased proteins in these pathways are shown in Supplementary Table 3. **C.** Proteomics detection of glycolysis enzyme proteins. Results are expressed as mean ± SEM. The p values were calculated with t-test. “\*”, <0.05, “\*\*\*”, <0.01; “\*\*\*\*”, <0.001; “\*\*\*\*\*”, <0.0001.

A

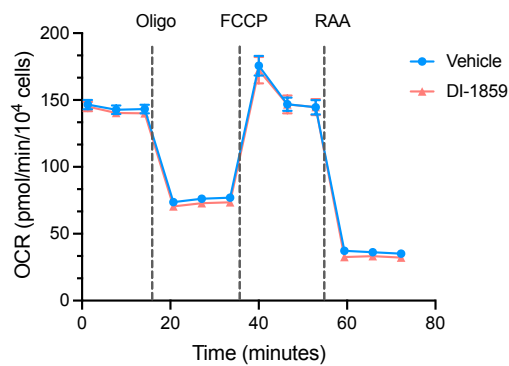

B

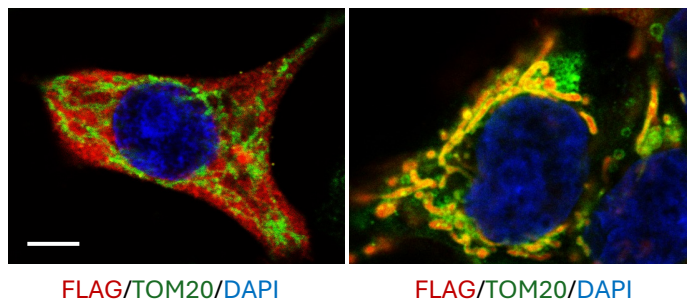

C

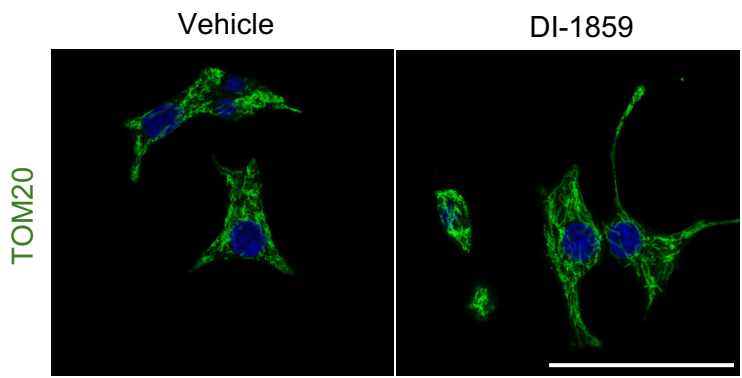

D

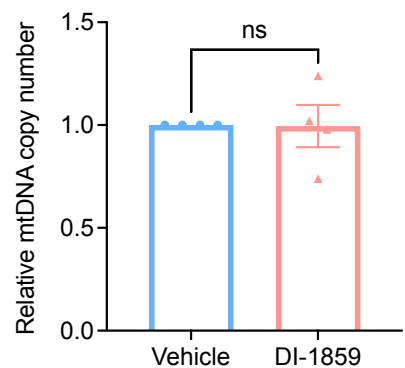

**Supplementary Figure 3. DI-1859 treatment does not alter mitochondria bioenergetic function or abundance in INS-1 832/13 cells.** **A.** Seahorse Flux assay of mitochondrial respiration in INS-1 832/13 cells. On the day of measurements, cells were pre-treated with Vehicle or 100 nM DI-1859 for 2 h in DMEM-XF (2.5 mM Glucose). Cells were then switched to Seahorse bicarbonate-free RPMI-XF (Aglient 103576-100, supplemented with 12.5 mM glucose, 2 mM L-glutamine, 1 mM sodium pyruvate, pH 7.4), with Vehicle or 100 nM DI-1859 treatment, and equilibrated at 37 °C non-CO<sub>2</sub> incubator for 1 h. Oxygen consumption rate (OCR) was monitored at basal state and after sequential injection of the mitochondrial compounds oligomycin (2.5 μM), FCCP (0.5 μM) and Rotenone/antimycin A (0.5 μM) that induce mitochondrial stress. **B.** Left panel. INS-1 832/13 cells were infected with Ad-Cul3-FLAG. Confocal microscopy was used to detect the cellular localization of Cul3-FLAG with anti-FLAG antibody and TOM20 as a mitochondrial marker. Nuclei were stained with DAPI. Right panel. Confocal microscopy. HepG2 cells stably expressing a mitochondrial targeted protein mtSTH-FLAG (mitochondria-targeted soluble transhydrogenase from *E. coli*) was shown for comparison. Scale bar = 5 μm. **C.** INS-1 832/13 cells were treated with vehicle (DMSO) or DI-1859 (100 nM) for 3 hours. Immunofluorescence staining of TOM20 (green, Alexa Fluor 488) was performed and images were taken with a confocal microscope. Nuclei were stained with DAPI (blue). A representative image under each condition is shown. Scale bar = 50 μm. **D.** INS-1 832/13 cells were treated with vehicle (DMSO) or DI-1859 (100 nM) for 3 hours. Mitochondrial DNA copy number was measured by real-time PCR. The rat mtDNA primer set provided by the kit amplifies one of the most conserved regions on rat mtDNA, and the single copy reference (SCR) primer set amplifies a 100 bp region on rat chromosome 17 and serves as reference for data normalization. Relative mtDNA copy number was calculated using the $\Delta\Delta C_t$ method. The result is mean ± SEM of 4 independent experiments. The p value was calculated with unpaired t-test. ns. Not significant, $p > 0.05$.

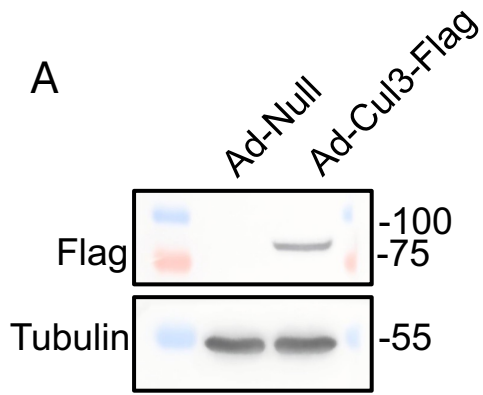

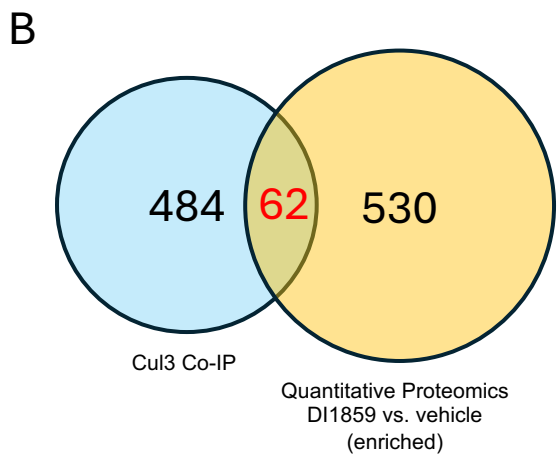

**Supplementary Figure 4. Co-IP proteomics detection of Cul3 associated proteins in INS-1 832/13 cells. A.** Western blot validation of Cul3-FLAG expression in INS-1 832/13 cells infected with Ad-Cul3-FLAG. The same samples were used for FLAG Co-IP proteomics. **B.** Venn diagram. Co-IP proteomics detection of Cul3 associated proteins that were also upregulated by DI-1859 in quantitative proteomics.
