## Supplementary Methods for "A selective Cullin 3 RING E3 ligase inhibitor attenuates hyperglycemia via dual insulin sensitizing and insulinotropic action"

**Reagents.** MLN4924, TAS4464 and DI-1859 were purchased from MedChemExpress, Inc. (Monmouth Junction, NJ). Insulin (Novolin, Cat. #: NDC 0169-1833-11) was purchased from Novo Nordisk, Inc. (Plainsboro Township, NJ). Cycloheximide and antibodies against Nedd8 (Cat. #: 2754, Lot. 2), Cul3 (Cat. #: 2759, Lot. 2), IRS1 (Cat. #: 2382, Lot. 10), p-AKT(S473) (Cat. #: 4060, Lot. 24), T-AKT (Cat. #: 4691, Lot.20), FLAG (Cat. #: 14793, Lot. 7. and #8146, Lot. 6), RhoA (Cat. #: 2117S, Lot. 6), MLC2 (Cat. #: 8505S, Lot. 6), p-MLC2 (Cat. #: 95777S, Lot. 3) were purchased from Cell Signaling Technology, Inc. (Danvers, MA). Antibodies against Cul1 (Cat. #: 32-2400, Lot.UJ297298), Cul2 (Cat. #: 51-1800, Lot.UA281866), and goat anti-rabbit Alexa Fluor 488 $2^{nd}$ antibody (Cat. #: A-11008) and goat anti-mouse Alexa-Fluor 594 $2^{nd}$ antibody (Cat. #: A-11005) were purchased from Invitrogen (Carlsbad, CA). Antibodies against Cul4B (Cat. #: 12916-I-AP) and TOM20 (Cat. #: 11802-1-AP) were purchased from Proteintech Group, Inc. (Rosemont, IL). Streptozocin (STZ, Cat. #: S0130), 2-hydroxypropyl-β-cyclodextrin (Cat. #: 332607), and Anti-FLAG magnetic beads were purchased from Sigma Aldrich (St. Louis, MO). Antibodies against Actin (Cat. #: Ab3280) was purchased from Abcam (Cambridge, MA). Insulin rodent (mouse/rat) chemiluminescence ELISA kit (Cat. #: 80-INSMR-CH10) and human insulin ELISA kit (Cat. #: 80-INSHU-E01.1) were purchased from ALPCO Diagnostics (Salem, NH). Lipofectamine RNAiMAX reagent was purchased from ThermoFisher Scientific (Waltham, MA). Glucose Uptake-Glo Assay kit (Cat. #: K1341) was purchased from Promega (Madison, WI). Mouse GLP-1 ELISA kit (Cat. #: 81508; lot: 62101202) and C-peptide mouse ELISA kit (Cat. #: #90050) were purchased from Crystal Chem (Elk Grove Village, IL). RhoA inhibitor (C3, Cat. #: CT04) was purchased from Cytoskeleton, Inc. (Denver, CO). Latrunculin A (LAT-A, Cat. #: 10010630) was purchased from Cayman Chemical (Ann Arbor, MI). Phalloidin-California Red Conjugate (Cat. #: sc-499440) was purchased from Santa Cruz Biotechnology, Inc. (Dallas, TX). The siGENOME SMARTpool siCul3 against rat Cul3 and siControl were purchased from Horizon Discovery (Lafayette, CO).

**Co-immunoprecipitation-proteomics.** The immunoprecipitated samples in Laemmli buffer were run on SDS-PAGE followed by trypsin in-gel digestion. Each digested sample was analyzed by nano LC-MS/MS with a Waters M-Class HPLC system interfaced to a ThermoFisher Exploris 480 mass spectrometer. Peptides were loaded on a trapping column and eluted over a 75 µm analytical column at 350nL/min; both columns were packed with Luna C18 resin (Phenomenex). The mass spectrometer was operated in data-dependent mode, with the Orbitrap operating at 60,000 FWHM and 15,000 FWHM for MS and MS/MS respectively. APD was enabled and the instrument was run with a 3s cycle for MS and MS/MS. Five hrs of instrument time was used for the analysis of each sample. Data were searched using a local copy of Mascot (Matrix Science). Mascot DAT files were parsed into Scaffold (Proteome Software) for validation, filtering and to create a non-redundant list per sample. Data were filtered using at 1% protein and peptide FDR and requiring at least two unique peptides per protein. An interacting protein is assigned based on the following criteria: a protein had at least 5 Spectral counts (SpC) in the IP sample and was not detected in the NC sample; or a protein was detected with a 4-fold or more SpC in IP sample over NC sample.

**Quantitative Proteomics**. Following the same condition under which insulin secretion was studied, INS-1 832/13 cells were cultured to ~90% confluence in RMPI1640 medium (12.5 mM glucose, supplemented with 2 mM L-glutamine, 1 mM sodium pyruvate, 10 mM HEPES, pH 7.4, 0.05 mM β-mercaptoethanol, penicillin–streptomycin, and 10% FBS). Cells were then cultured in DMEM (Gibco, Cat. #: A14430-01, supplemented with 2.5 mM glucose, 2 mM L-glutamine, 1 mM sodium pyruvate, 10 mM HEPES, pH-7.4, 0.05 mM β-mercaptoethanol, 1%

penicillin–streptomycin) overnight. The next morning, cells were treated with DI-1859 (100 nM) or vehicle (DMSO) for 3 hours. Cells were washed in cold PBS 3 times and lysed in 1X RIPA buffer and heated at 70 °C for 15 minutes. Protein concentration was measured and 200 mg protein was mixed with 14 pmol of horse serum albumin and precipitated overnight with 4 volumes of cold acetone. The precipitated protein was reconstituted in 100 ml Laemmli buffer to a final concentration of 1 mg/ml, and 20 ml of this sample was run on an SDS-PAGE gel. Each lane was cut as a sample, chopped into smaller pieces, washed, reduced, alkylated, and digested with 1 µg trypsin overnight at room temperature.  Peptides were extracted from the gel in 50% acetonitrile. The extracts were dried by a SpeedVac and reconstituted in 200 µl 1% acetic acid for analysis. An Orbitrap Exploris 480 Mass Spectrometer (ThermoFisher Scientific) was used in the data-independent acquisition (DIA) mode with a 10m/z window working from m/z 350 to 950.  The orbitrap was operated at a resolution of 15,000.  A full scan spectrum at a resolution of 35,000 was acquired each cycle. Data were analyzed using the DIA-NN program. The results were normalized to the horse serum albumin internal standard. Unpaired Student's t-test was used for detecting significant changes ($p <0.05$, fold change cutoff: 1.2 or -0.8). Differentially regulated proteins were analyzed using Ingenuity Pathway Analysis (IPA).

**Detection of glycolysis intermediate metabolites in U-$C_{13}$-glucose metabolic tracing through glycolysis**. Briefly, a 50 µL aliquot of the cell lysate sample or each of serially diluted standard solutions of glucose, ribose-5P, glycerylaldehyde-3P, glucose-6P was mixed with 100 µL of 25-mM AEC solution, 50 µL of 50 mM $NaCBH3$ solution and 20 µL of acetic acid. The mixtures were allowed to react at 60°C for 70 min. After reaction, 300 µL of water and 300 µL of chloroform were added. After vortex-mixing for 15 s and centrifugation for 5 minutes, 10 µL aliquots of the resultant solutions were injected into a PFP column (2.1 x 150 mm, 1.7 µm) to run

UPLCMRM/MS on an Agilent 1290 UHPLC system coupled to an Agilent 6495B QQQ instrument with positive-ion detection. For detection of other metabolites, serially diluted standard solutions of all the targeted metabolites were prepared in 75% methanolacetonitrile (1:1). Twenty µL of the clear supernatant of each sample or 20 µL of each standard solution was mixed with 180 µL of water. Ten µL of each sample solution or each standard solution was injected into a C18 UPLC column to run LC-MRM/MS with negative ion detection on a Waters Acquity UPLC system coupled to a Sciex QTRAP 6500 Plus MS instrument, with the use of a tributylamine buffer and acetonitrile as the mobile phase for gradient elution. Peak areas of U-$C_{12}$ forms and $C_{13}$ forms of detected metabolites were calculated.

**Confocal microscopy.** After treatments, INS-1 832/13 cells were fixed with 4% paraformaldehyde in PBS and permeabilization was performed using 0.1% Triton X-100 in PBS. Cells were blocked with 5% normal goat serum in PBS containing 0.1% Triton X-100 for 1 hour at room temperature. Cells were then incubated overnight at 4°C with a primary antibody against TOM20 (Proteintech Group, Inc. Cat. #: 11802-1-AP, lot #. 00169998) diluted 1:200 in blocking buffer or FLAG (Cell Signaling Technology, Inc. Cat. #8146, Lot. X). The following day, cells were washed three times with PBS and incubated for 1 hour at room temperature with an Alexa Fluor 488-conjugated secondary antibody (Goat anti-Mouse IgG (H+L) Cross-Adsorbed Secondary Antibody, Alexa Fluor™ 488, (ThermoFisher Scientific, Cat. #: A-11001, 1:1000 dilution) or Goat anti-Mouse IgG (H+L) Cross-Adsorbed Secondary Antibody, Alexa Fluor™ 594 (ThermoFisher Scientific, Cat. #: A-11005, 1:1000 dilution) in PBS. After final washes, coverslips were mounted using a DAPI-containing mounting medium (Cat. #: H-1200, Vector laboratories, Inc. Newark, CA). Fluorescence images were acquired using a Leica SP8 confocal microscope with consistent acquisition settings across samples.
