## Supplementary Table 1 for "A selective Cullin 3 RING E3 ligase inhibitor attenuates hyperglycemia via dual insulin sensitizing and insulinotropic action"

### Supplementary Table 1. Cul3 association with CRL3 core components

| Identified Proteins | Rank | NC (SpC) | Cul3 IP (SpC) |
|---|---|---|---|
| **Cullin-3 GN=Cul3** | 1 | 9 | 3192 |
| E3 ubiquitin-protein ligase RBX1 GN=Rbx1 | 169 | 0 | 16 |
| BTB (POZ) domain containing 1 GN=Btbd1 | 51 | 0 | 64 |
| BTB domain containing 10 GN=Btbd10 | 61 | 0 | 52 |
| BTB domain containing 2 GN=Btbd2 | 27 | 0 | 114 |
| BTB domain containing 3 GN=Btbd3 | 54 | 0 | 59 |
| BTB domain containing 6 GN=Btbd6 | 140 | 0 | 19 |
| **BTB/POZ domain-containing adapter for CUL3-mediated RhoA degradation protein 1 GN=Bacurd1** | 29 | 0 | 111 |
| **BTB/POZ domain-containing adapter for CUL3-mediated RhoA degradation protein 2 GN=Bacurd2** | 16 | 0 | 160 |
| **BTB/POZ domain-containing adapter for CUL3-mediated RhoA degradation protein 3 GN=Bacurd3** | 9 | 0 | 187 |
| BTB/POZ domain-containing protein 7 GN=Btbd7 | 192 | 0 | 14 |
| BTB/POZ domain-containing protein 8 GN=Btbd8 | 37 | 7 | 86 |
| BTB/POZ domain-containing protein 9 GN=Btbd9 | 14 | 0 | 173 |
| BTB/POZ domain-containing protein KCTD3 GN=Kctd3 | 18 | 0 | 154 |
| BTB/POZ domain-containing protein KCTD7 GN=Kctd7 | 73 | 0 | 46 |
| BTB/POZ domain-containing protein KCTD9 GN=Kctd9 | 58 | 0 | 55 |
| BTB/POZ domain-containing protein KCTD21 GN=Kctd21 | 81 | 0 | 37 |
| BTB/POZ domain-containing protein KCTD18 GN=Kctd18 | 97 | 0 | 28 |
| Kelch repeat and BTB domain containing 2 GN=Kbtbd2 | 26 | 0 | 114 |
| Kelch repeat and BTB domain containing 4 GN=Kbtbd4 | 21 | 0 | 137 |
| Kelch repeat and BTB domain containing 7 GN=Kbtbd7 | 44 | 0 | 77 |
| Kelch repeat and BTB domain containing 8 GN=Kbtbd8 | 63 | 0 | 49 |
| Kelch-like ECH-associated protein 1 GN=Keap1 | 24 | 3 | 115 |
| Kelch-like family member 13 GN=Klhl13 | 10 | 0 | 187 |
| Kelch-like family member 18 GN=Klhl18 | 132 | 0 | 20 |
| Kelch-like family member 23 GN=Klhl23 | 34 | 0 | 92 |
| Kelch-like family member 26 GN=Klhl26 | 19 | 0 | 149 |
| Kelch-like family member 28 GN=Klhl28 | 466 | 0 | 6 |
| Kelch-like family member 32 GN=Klhl32 | 56 | 0 | 57 |
| Kelch-like family member 5 GN=Klhl5 | 150 | 0 | 18 |
| Kelch-like family member 8 GN=Klhl8 | 32 | 0 | 106 |
| Kelch-like family member 9 GN=Klhl9 | 3 | 0 | 374 |
| Kelch-like protein 11 GN=Klhl11 | 126 | 0 | 21 |
| Kelch-like protein 12 GN=Klhl12 | 50 | 0 | 64 |
| Kelch-like protein 15 GN=Klhl15 | 60 | 0 | 53 |
| Kelch-like protein 17 GN=Klhl17 | 506 | 0 | 5 |
| Kelch-like protein 2 GN=Klhl2 | 17 | 0 | 155 |
| Kelch-like protein 20 GN=Klhl20 | 42 | 0 | 80 |
| Kelch-like protein 21 GN=Klhl21 | 68 | 0 | 47 |
| Kelch-like protein 22 GN=Klhl22 | 4 | 4 | 364 |
| Kelch-like protein 24 GN=Klhl24 | 25 | 0 | 114 |
| Kelch-like protein 25 GN=Klhl25 | 12 | 0 | 178 |
| Kelch-like protein 3 GN=Klhl3 | 349 | 0 | 7 |
| Kelch-like protein 42 GN=Klhl42 | 23 | 0 | 124 |
| Kelch-like protein 7 GN=Klhl7 | 7 | 0 | 217 |
| COP9 signalosome complex subunit 2 GN=Cops2 | 11 | 3 | 186 |
| COP9 signalosome complex subunit 3 GN=Cops3 | 46 | 2 | 71 |
| COP9 signalosome complex subunit 4 GN=Cops4 | 30 | 5 | 110 |
| COP9 signalosome complex subunit 5 GN=Cops5 | 39 | 0 | 85 |
| COP9 signalosome subunit 6 GN=Cops6 | 36 | 2 | 88 |
| COP9 signalosome subunit 7A GN=Cops7a | 90 | 5 | 31 |
| COP9 signalosome subunit 7B GN=Cops7b | 59 | 0 | 54 |
| COP9 signalosome complex subunit 8 GN=Cops8 | 156 | 0 | 17 |

**Note**: NC: negative control for IP; SpC: Spectral count. Rank: Abundance of SpC in Cul3 IP sample.
