## Supplementary Table 2 for "A selective Cullin 3 RING E3 ligase inhibitor attenuates hyperglycemia via dual insulin sensitizing and insulinotropic action"

**Supplementary Table 2. Cul3-associated proteins that are upregulated by DI-1859**

| Symbol | Protein name |
|---|---|
| **AIP** | AH receptor-interacting protein |
| **AK1** | Adenylate kinase isoenzyme 1 |
| **ARG1** | Arginase-1 |
| **ARPC3** | Actin-related protein 2/3 complex subunit 3 |
| **ASAH1** | Acid ceramidase |
| **ATP5PD** | ATP synthase subunit d, mitochondrial |
| **BAG2** | BAG cochaperone 2 |
| **CDIPT** | CDP-diacylglycerol--inositol 3-phosphatidyltransferase |
| **CDK4** | Cyclin-dependent kinase 4 |
| **CLTB** | Clathrin light chain B |
| **CNP** | 2',3'-cyclic-nucleotide 3'-phosphodiesterase |
| **COPE** | Coatomer subunit epsilon |
| **COPS3** | COP9 signalosome complex subunit 3 |
| **COPS7B** | COP9 signalosome subunit 7B |
| **CPLX2** | Complexin-2 |
| **CR1L** | Complement component receptor 1-like protein |
| **CRKL** | Crk-like protein |
| **CSTF2** | Cleavage stimulation factor, 3' pre-RNA subunit 2 |
| **CUSTOS** | Protein CUSTOS |
| **DUSP3** | Dual specificity protein phosphatase |
| **ECSIT** | Evolutionarily conserved signaling intermediate in Toll pathway, mitochondrial |
| **GORASP2** | Golgi reassembly-stacking protein 2 |
| **HAPLN4** | Hyaluronan and proteoglycan link protein 4 |
| **HDGF** | Hepatoma-derived growth factor |
| **HIBADH** | 3-hydroxyisobutyrate dehydrogenase, mitochondrial |
| **LIG3** | DNA ligase |
| **LMAN2** | Lectin, mannose-binding 2 |
| **MANF** | Mesencephalic astrocyte-derived neurotrophic factor |
| **MBNL2** | Muscleblind-like protein 2 |
| **MCU** | Calcium uniporter protein |
| **MINPP1** | Multiple inositol polyphosphate phosphatase 1 |
| **MLXIPL** | Carbohydrate-responsive element-binding protein |
| **MRPL3** | Large ribosomal subunit protein uL3m |
| **MRPS9** | Mitochondrial ribosomal protein S9 |
| **NAA50** | N(alpha)-acetyltransferase 50, NatE catalytic subunit |
| **NACC1** | Nucleus accumbens-associated protein 1 |

| | |
|---|---|
| **NAGA** | Alpha-N-acetylgalactosaminidase |
| **NISCH** | Nischarin |
| **NUBP1** | Cytosolic Fe-S cluster assembly factor NUBP1 |
| **PCSK2** | Neuroendocrine convertase 2 |
| **PDP2** | [Pyruvate dehydrogenase [acetyl-transferring]]-phosphatase 2, mitochondrial |
| **PGP** | Glycerol-3-phosphate phosphatase |
| **PIP4K2C** | Phosphatidylinositol 5-phosphate 4-kinase type-2 gamma |
| **PITPNA** | Phosphatidylinositol transfer protein alpha isoform |
| **PPP2CA** | Serine/threonine-protein phosphatase 2A catalytic subunit alpha isoform |
| **PPT1** | Palmitoyl-protein thioesterase 1 |
| **PSMB2** | Proteasome subunit beta type-2 |
| **PSME2** | Proteasome activator complex subunit 2 |
| **PSMF1** | Proteasome inhibitor PI31 subunit |
| **PUS1** | Pseudouridylate synthase 1 homolog |
| **RAB43** | Ras-related protein Rab-43 |
| **RAP1B** | Ras-related protein Rap-1b |
| **RPRD1B** | Regulation of nuclear pre-mRNA domain-containing protein |
| **SAE1** | SUMO-activating enzyme subunit 1 |
| **SEZ6L2** | Seizure related 6 homolog like 2 |
| **TEX2** | Testis expressed 2 |
| **TMED10** | Transmembrane emp24 domain-containing protein 10 |
| **UBE2Z** | Ubiquitin-conjugating enzyme E2 Z |
| **UBXN1** | UBX domain-containing protein 1 |
| **UCK1** | Uridine-cytidine kinase |
| **VAT1L** | Vesicle amine transport 1-like |
| **WBP2** | WW domain-binding protein 2 |
