## Supplementary Table 3 for "A selective Cullin 3 RING E3 ligase inhibitor attenuates hyperglycemia via dual insulin sensitizing and insulinotropic action"

**Supplementary Table 3. Upregulated pathways by DI-1859**

| Ingenuity Canonical Pathways | -log$^{(p)}$ | Ratio | z-score | Upregulated Proteins |
|---|---|---|---|---|
| RHO GTPases Activate Formins | 7.28 | 0.114 | 4 | BUB3,CDC42,CENPU,DYNLL1,DYNLL2, MAPRE1,NUP85,PFN1,PFN2,PPP2CA, PPP2R5C,RAC1, RHOA,TUBAL3,TUBB4A,ZWINT |
| RHO GTPases activate PKNs | 4.65 | 0.214 | 2.449 | MYL6,PPP1CB,RAC1,RHOA,YWHAG,YWHAZ |
| RHO GTPases activate KTN1 | 4.23 | 0.364 | 2 | CDC42,RAC1,RHOA,RHOG |
| RHO GTPases activate PAKs | 4.19 | 0.238 | 1.342 | CALM1,CDC42,MYL6,PPP1CB,RAC1 |
| RHO GTPases activate IQGAPs | 3.27 | 0.156 | 1.342 | CALM1,CDC42,RAC1,TUBAL3,TUBB4A |
| RHO GTPases activate CIT | 3.22 | 0.211 | 2 | MYL6,PPP1CB,RAC1,RHOA |
| RHO GTPases Activate ROCKs | 3.22 | 0.211 | 2 | CFL1,MYL6,PPP1CB,RHOA |
| RHOGDI Signaling | 2.11 | 0.05 | -2.53 | ARPC3,ARPC4,ARPC5L,CDC42,CFL1,GNG4, MYL6,PIP4K2C,RAC1,RHOA,RHOG |
| RHO GTPase cycle | 2.09 | 0.04 | 3.3 | BASP1,CDC42,DDRGK1,NISCH,NSFL1C, PGRMC2,RAC1,RACGAP1,RHOA,RHOG, SH3BP1,STIP1,TEX2,TPM3,TPM4,TXNL1,VAPB, VAV2 |
| RHOA Signaling | 3.52 | 0.0813 | 2.53 | ARPC3,ARPC4,ARPC5L,CFL1,MYL6,PFN1, PFN2,PIP4K2C,PPP1CB,RHOA |
| Regulation of Actin-based Motility by Rho | 6.15 | 0.117 | 2.887 | ARPC3,ARPC4,ARPC5L,CDC42,CFL1,MYL6, PFN1,PFN2,PIP4K2C,PPP1CB,RAC1,RHOA, RHOG |
| Integrin Signaling | 5.55 | 0.081 | 2.324 | ARF2,ARF4,ARF6,ARPC3,ARPC4,ARPC5L, CDC42,CRKL,NRAS,PARVB,PFN1,PFN2, PPP1CB,RAC1,RAP1B,RHOA,RHOG |
| EPH-Ephrin signaling | 5.34 | 0.118 | 3.317 | ARPC3,ARPC4,CDC42,CFL1,CLTA,CLTB, EFNB1,MYL6,RAC1,RHOA,VAV2 |
| Ephrin B Signaling | 4.62 | 0.123 | 1.633 | ACP1,CAP1,CDC42,CFL1,EFNB1,GNG4,RAC1, RHOA,VAV2 |
| Actin Cytoskeleton Signaling | 4.17 | 0.0661 | 2.84 | ARPC3,ARPC4,ARPC5L,CDC42,CFL1,CRKL, INS1,MYL6,NRAS,PFN1,PFN2,PPP1CB,RAC1, RAP1B, RHOA,VAV2 |
| Ephrin Receptor Signaling | 3.39 | 0.0644 | 3.317 | ACP1,ARPC3,ARPC4,ARPC5L,CDC42,CFL1, CRKL,EFNB1,GNG4,NRAS,RAC1,RAP1B,RHOA |
| Pyruvate metabolism | 4.13 | 0.143 | 2.236 | FAHD1,GLO1,GSTZ1,MPC2,PDHB,PDP2, VDAC1 |
| Glucose metabolism | 4.05 | 0.105 | 3 | ALDOA,GAPDH,NUP58,NUP62,NUP85,PGAM1, PGP,PPP2CA,TPI1 |
| Gluconeogenesis I | 3.41 | 0.167 | 2 | ALDOA,GAPDH,MDH1,MDH2,PGAM1 |
| NRF2-mediated Oxidative Stress Response | 3.61 | 0.059 | 2.646 | AKR1A1,CBR1,DNAJC9,ERP29,FTH1,GSTK1, GSTM7,GSTO1,GSTZ1,MGST1,NRAS,RAP1B, SOD1,SOD2,STIP1,UBE2K |
| KEAP1-NFE2L2 pathway | 2.8 | 0.0658 | 3.162 | CSNK2B,NME3,PSMA1,PSMB2,PSMD8,PSMD9, PSME1,PSME2,PSMF1,SKP1 |
| Glutathione-mediated Detoxification | 2.35 | 0.098 | 2 | GSTK1,GSTM7,GSTO1,GSTZ1,MGST1 |
