## Supplementary Table 4 for "A selective Cullin 3 RING E3 ligase inhibitor attenuates hyperglycemia via dual insulin sensitizing and insulinotropic action"

**Supplementary Table 4. RhoA associated proteins**

| Protein name | rank | NC (SpC) | RhoA IP (SpC) |
|---|---|---|---|
| **Ras homolog family member A GN=Rhoa** | 1 | 0 | 311 |
| **Rho GDP-dissociation inhibitor 1 GN=Arhgdia** | 2 | 4 | 166 |
| **Rap1 GTPase-GDP dissociation stimulator 1 GN=Rap1gds1** | 3 | 0 | 129 |
| **Actin, cytoplasmic 2 GN=Actg1** | 4 | 0 | 121 |
| **Rho guanine nucleotide exchange factor 1 GN=Arhgef1** | 5 | 7 | 35 |
| Ig kappa chain C region GN=Igkc | 6 | 0 | 19 |
| Matrin-3 GN=Matr3 | 7 | 4 | 18 |
| Ig gamma-2B chain C region GN=Igh-1a | 8 | 0 | 15 |
| Succinate dehydrogenase [ubiquinone] flavoprotein subunit, mitochondrial GN=Sdha | 9 | 4 | 15 |
| Small ribosomal subunit protein uS14 GN=Rps29 | 10 | 3 | 11 |
| Rho guanine nucleotide exchange factor 12 GN=Arhgef12 | 11 | 3 | 11 |
| Nucleolar and coiled-body phosphoprotein 1 GN=Nolc1 | 12 | 3 | 11 |
| Signal recognition particle receptor subunit beta GN=Tf | 13 | 2 | 10 |
| Protein disulfide-isomerase A2 GN=Pdia2 | 14 | 0 | 8 |
| N(alpha)-acetyltransferase 16, NatA auxiliary subunit GN=Naa16 | 15 | 0 | 8 |
| Ubiquitin conjugating enzyme E2 V1 GN=Ube2v1 | 16 | 2 | 8 |
| Dishevelled associated activator of morphogenesis 1 GN=Daam1 | 17 | 0 | 7 |
| Cold-inducible RNA-binding protein GN=Cirbp | 18 | 2 | 7 |
| Ubiquitin carboxyl-terminal hydrolase 30 GN=Usp30 | 19 | 0 | 6 |
| NHP2-like protein 1 GN=Snu13 | 20 | 0 | 6 |
| ATP-dependent RNA helicase DDX39A GN=Ddx39a | 21 | 0 | 6 |
| Neuronal guanine nucleotide exchange factor GN=Ngef | 22 | 0 | 6 |
| Signal peptidase complex subunit 2 GN=Spcs2 | 23 | 0 | 6 |
| Kelch-like protein 22 GN=Klhl22 | 24 | 0 | 5 |
| Histone H3.3 GN=H3-3b | 25 | 0 | 5 |
| F-box only protein 30 GN=Fbxo30 | 26 | 0 | 5 |
| Zinc phosphodiesterase ELAC protein 2 GN=Elac2 | 27 | 0 | 5 |
| WD repeat domain 43 GN=Wdr43 | 28 | 0 | 5 |
| Keratin 31 GN=Krt31 | 29 | 0 | 5 |
| Ribosomal L1 domain-containing protein 1 GN=Rsl1d1 | 30 | 0 | 5 |

**Note**: NC: negative control for IP; SpC: Spectral count. Rank: Abundance of SpC in RhoA IP sample.
