## Supplementary Table 5 for "A selective Cullin 3 RING E3 ligase inhibitor attenuates hyperglycemia via dual insulin sensitizing and insulinotropic action"

### Supplementary Table 5. Cul3 association with cytoskeleton proteins

| Identified Proteins | NC (SpC) | Cul3 IP (SpC) |
|---|---|---|
| **Core cytoskeletal proteins** | | |
| Tubulin alpha-1A chain GN=Tuba1a | 70 | 472 |
| F-actin-capping protein subunit alpha-2 GN=Capza2 | 3 | 17 |
| Keratin, type II cytoskeletal 7 GN=Krt7 | 0 | 7 |
| Septin-7 GN=Septin | 0 | 10 |
| Septin-5 GN=Septin5 | 0 | 5 |
| **Actin-associated / cytoskeleton regulators** | | |
| Actin-related protein 2/3 complex subunit 3 GN=Arpc3 | 0 | 8 |
| Coronin-1A OX=10116 GN=Coro1a PE=1 SV=3 | 0 | 7 |
| Shroom family member 2 GN=Shroom2 | 0 | 7 |
| Serine/threonine-protein kinase MRCK gamma GN=Cdc42bpg | 0 | 5 |
| Formin-binding protein 1-like GN=Fnbp1l | 0 | 6 |
| Ras suppressor protein 1 GN=Rsu1 | 0 | 5 |
| LIM and senescent cell antigen-like-containing domain protein GN=Lims1 | 0 | 5 |
| Tensin 3 GN=Tns3 | 0 | 5 |
| PTPRF interacting protein alpha 1 GN=Ppfia1 | 0 | 13 |
| PTPRF interacting protein alpha 3 GN=Ppfia3 | 0 | 9 |
| Huntingtin-associated protein 1 GN=Hap1 | 0 | 10 |
| **Microtubule-associated proteins** | | |
| TPX2, microtubule nucleation factor GN=Tpx2 | 0 | 16 |
| Protein regulator of cytokinesis 1 GN=Prc1 | 4 | 24 |
| Centrosomal protein of 170 kDa GN=Cep170 | 3 | 104 |
| Centrosomal protein 170B GN=Cep170b | 0 | 43 |
| Gamma-tubulin complex component 6 GN=Tubgcp6 | 0 | 7 |
| G-2 and S-phase expressed 1 GN=Gtse1 | 0 | 10 |
| MAP7 domain-containing protein 1 GN=Map7d1 | 0 | 8 |
| Calmodulin-regulated spectrin-associated protein 2 GN=Camsap2 | 0 | 13 |
| CAP-GLY domain containing linker protein 1 GN=Clip1 | 0 | 6 |
| Hyaluronan-mediated motility receptor GN=Hmmr | 0 | 12 |
| **Note**: NC: negative control for IP; SpC: Spectral count | | |
